# Reconstructing the history of variation in effective population size along phylogenies

**DOI:** 10.1101/793059

**Authors:** Mathieu Brevet, Nicolas Lartillot

## Abstract

The nearly-neutral theory predicts specific relations between effective population size (*N_e_*) and patterns of divergence and polymorphism, which depend on the shape of the distribution of fitness effects (DFE) of new mutations. However, testing these relations is not straightforward, owing to the difficulty in estimating *N_e_*. Here, we introduce an integrative framework allowing for an explicit reconstruction of the phylogenetic history of *N_e_*, thus leading to a quantitative test of the nearly-neutral theory and an estimation of the allometric scaling of the ratios of non-synonymous over synonymous polymorphism (*π_N_ /π_S_*) and divergence (*dN/dS*) with respect to *N_e_*. As an illustration, we applied our method to primates, for which the nearly-neutral predictions were mostly verified. Under a purely nearly-neutral model with a constant DFE across species, we find that the variation in *π_N_ /π_S_* and *dN/dS* as a function of *N_e_* is too large to be compatible with current estimates of the DFE based on site frequency spectra. The reconstructed history of *N_e_* shows a ten-fold variation across primates. The mutation rate per generation *u*, also reconstructed over the tree by the method, varies over a three-fold range and is negatively correlated with *N_e_*. As a result of these opposing trends for *N_e_* and *u*, variation in *π_S_* is intermediate, primarily driven by *N_e_* but substantially influenced by *u*. Altogether, our integrative framework provides a quantitative assessment of the role of *N_e_* and *u* in modulating patterns of genetic variation, while giving a synthetic picture of their history over the clade.

**Significance statement:** Natural selection tends to increase the frequency of mutants of higher fitness and to eliminate less fit genetic variants. However, chance events over the life of the individuals in the population are susceptible to introduce deviations from these trends, which are expected to have a stronger impact in smaller populations. In the long-term, these fluctuations, called random drift, can lead to the accumulation of mildly deleterious mutations in the genomes of living species, and for that reason, the effective population size (usually denoted *N_e_*, and which captures the relative strength of drift, relative to selection) has been proposed as a major determinant of the evolution of genome architecture and content. A proper quantitative test of this hypothesis, however, is hampered by the fact that *N_e_* is difficult to estimate in practice. Here, we propose a Bayesian integrative approach for reconstructing the broad-scale variation in *N_e_* across an entire phylogeny, which in turns allows for quantifying how *N_e_* correlates with life history traits and with various measures of genetic diversity and selection strength, between and within species. We apply this approach to the phylogeny of primates, and observe that selection is indeed less efficient in primates characterized by smaller effective population sizes.

## Introduction

Effective population size (*N_e_*) is a central parameter in population genetics and in molecular evolution, impacting both genetic diversity and the strength of selection (Charlesworth, 2009; Leffler *et al.*, 2012). The influence of *N_e_* on diversity reflects the fact that larger populations can store more genetic variation, while the second aspect, efficacy of selection, is driven by the link between *N_e_* and genetic drift: the lower the *N_e_*, the more genetic evolution is influenced by the random sampling of individuals over generations. As a result, long-term trends in *N_e_* are expected to have an important impact on genome evolution (Lynch *et al.*, 2011) and, more generally, on the relative contribution of adaptive and non-adaptive forces in shaping macro-evolutionary patterns.

The nearly-neutral theory proposes a simple conceptual framework for formalizing the role of selection and drift on genetic sequences. According to this theory, genetic sequences are mostly under purifying selection; deleterious mutation are eliminated by selection, whereas neutral and nearly-neutral mutations are subject to genetic drift and can therefore segregate and reach fixation. The inverse of *N_e_* defines the selection threshold under which genetic drift dominates. This results in specific quantitative relations between *N_e_* and key molecular parameters (Ohta, 1995). In particular, species with small *N_e_* are expected to have a higher ratio of nonsynonymous (*d_N_*) to synonymous (*d_S_*) substitution rates and a higher ratio of nonsynonymous (*π_N_*) to synonymous (*π_S_*) nucleotide diversity. Under certain assumptions, these two ratios are linked to *N_e_* through allometric functions in which the scaling coefficient is directly related to the shape of the distribution of fitness effects (DFE) (Kimura, 1979; Welch *et al.*, 2008).

The empirical test of these predictions raises the problem that *N_e_* is difficult to measure directly in practice. In principle, *N_e_* could be estimated through demographic and census data. However, the relation between census and effective population size is far from straightforward. Consequently, many studies which have tried to test nearly-neutral theory have used proxies indirectly linked to *N_e_*. In particular, life history traits (LHT, essentially body mass or maximum longevity) are expected to correlate negatively with *N_e_* (Waples *et al.*, 2013). As a result, *d_N_ /d_S_* or *π_N_ /π_S_* are predicted to correlate positively with LHT. This has been tested, leading to various outcomes, with both positive and negative results (Eyre-Walker *et al.*, 2002; Popadin *et al.*, 2007; Nikolaev *et al.*, 2007; Lartillot, 2013; Nabholz *et al.*, 2013; Romiguier *et al.*, 2014; Figuet *et al.*, 2016).

More direct estimations of *N_e_* can be obtained from *π_S_* since, in accordance with coalescent theory, *π_S_* = 4*N_e_u* (with *u* referring to the mutation rate per site per generation). Thus, one would predict a negative correlation of *d_N_ /d_S_*, *π_N_ /π_S_* and LHT with *π_S_*. Such predictions have been tested, and generally verified, in several previous studies (Piganeau & Eyre-Walker, 2009; Romiguier *et al.*, 2014; Figuet *et al.*, 2016; Galtier, 2016; Chen *et al.*, 2017; James *et al.*, 2017). However, these more specific tests of the nearly-neutral theory are only qualitative, at least in their current form, in which *N_e_* is indirectly accessed through *π_S_* without any attempt to correct for the confounding effect of the mutation rate *u* and its variation across species.

## New Approaches

In this study, we aim to solve this problem by using a Bayesian integrative approach, in which the joint evolutionary history of a set of molecular and phenotypic traits is explicitely reconstructed along a phylogeny. This method has previously been used to test the predictions of the nearly-neutral theory via indirect proxies of *N_e_* (Lartillot, 2013; Nabholz *et al.*, 2013). Here, we propose an elaboration on this approach, in which the variation in the mutation rate per generation *u* is globally reconstructed over the phylogeny by combining the relaxed molecular clock of the model with data about generation times. This in turns allows us to tease out *N_e_* and *u* from the *π_S_* estimates obtained in extant species, thus leading to a complete reconstruction of the phylogenetic history of *N_e_* and of its scaling relations with others traits such as *d_N_ /d_S_* or *π_N_ /π_S_*. Using this reconstruction, we can conduct a proper quantitative test of some of the predictions of the nearly-neutral theory and then compare our findings with independent knowledge previously derived from the analysis of site frequency spectra. The approach requires a multiple sequence alignment across a group of species, together with polymorphism data, ideally averaged over many loci to stabilize the estimates, as well as data about life-history traits in extant species and fossil calibrations. Here, we apply it to previously published phylogenetic and transcriptome data in primates (Perelman *et al.*, 2011; Perry *et al.*, 2012).

## Results

### Empirical phylogenetic correlation analysis

We first conducted a general correlation analysis between *dS*, *dN/dS*, *π_S_*, *π_N_ /π_S_* and life-history traits (LHTs). Our analysis relies on a multiple sequence alignment of 54 coding genes in 61 primates (adapted from Perelman *et al.*, 2011), combined with data about LHTs taken from the literature (de Magalhaes & Costa, 2009; Besenbacher *et al.*, 2019) and polymorphism data from 10 primates species (obtained from Romiguier *et al.*, 2014; Figuet *et al.*, 2016; Perry *et al.*, 2012). Of note, these estimates of *π_S_* and *π_N_ /π_S_* are based on the global transcriptome data, thus not restricted to the 54 coding genes represented in the multiple sequence alignment (see methods for details, Data subsection).

In a first step, these data were analysed using a previously introduced Bayesian integrative model (Coevol, Lartillot & Poujol, 2011). This model is an adaptation of the classical comparative method based on the principle of phylogenetically independent contrasts (Felsenstein, 1985) to the problem of estimating the correlation between quantitative traits (such as LHT, *π_S_* and *π_N_ /π_S_*) and substitution rates (such as *dS* and *dN/dS*). The joint evolutionary process followed by all these variables is assumed to be multivariate Brownian:

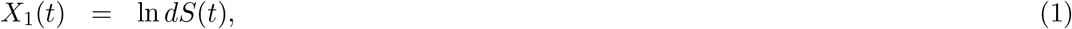

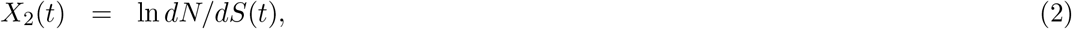

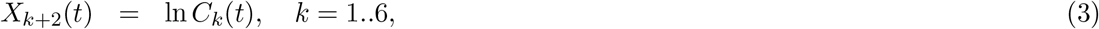

where *C_k_*(*t*) are the instant values of the 6 traits (body mass, age at sexual maturity, longevity, generation time *π_S_*, *π_N_ /π_S_*). This process is parameterized by a 8 *×* 8 variance-covariance matrix Σ, which captures the correlation structure between rates and traits. The model is conditioned on sequence and trait data, using Markov Chain Monte-Carlo (MCMC). Then, the marginal posterior distribution on Σ is used to derive measures of the strength and the slopes of the correlations between rates and traits, using standard covariance analysis and in a way that automatically accounts for phylogenetic inertia. In the following, this Brownian model is referred to as the *phenomenological* model, because it does not entail any specific assumption about the population-genetic relations that might exist between *dN/dS*, *π_N_ /π_S_* and *N_e_*.

Based on this first analysis, neither *π_N_ /π_S_* nor *dN/dS* appear to correlate with LHTs (Table 1), except for a positive correlation between *d_N_ /d_S_* and longevity (correlation coefficient *r* = 0.49). On the other hand, the correlations among molecular quantities are globally in agreement with the nearly-neutral predictions, although with rather unequal statistical support. Most notably, *π_N_ /π_S_* shows a clear negative correlation with *π_S_* (*r* = *−*0.73). As for *dN/dS*, it also shows a negative correlation with *π_S_* (*r* = *−*0.50), although with very marginal support (posterior probability of a negative correlation *pp <* 0.95). The two variables, *dN/dS* and *π_N_ /π_S_* are also positively correlated with each other (*r* = 0.42), but again, with marginal support. The weaker correlations observed for *dN/dS*, compared to the more robust correlation between *π_N_ /π_S_* and *π_S_*, could be due either to the presence of a minor fraction of adaptive substitutions or, alternatively, to a discrepancy between the short-term demographic effects reflected in both *π_S_* and *π_N_ /π_S_* and long-term trends captured by *dN/dS*.

**Table 1.** Correlation coefficients between *dS*, *dN/dS*, *π_S_* and *π_N_ /π_S_*, life-history traits and *N_e_*.

| | $dS$ | $dN/dS$ | Mat. | Mass | Long. | $\pi_S$ | $\pi_N/\pi_S$ | Gen. | $u$ | $N_e$ |
| --- | --- | --- | --- | --- | --- | --- | --- | --- | --- | --- |
| $dS$ | | 0.24 | -0.38 | -0.64** | -0.27 | -0.60 | 0.45 | -0.42 | 0.72** | -0.72** |
| $dN/dS$ | | | 0.15 | 0.08 | 0.49 | -0.50 | 0.42 | 0.42 | 0.58* | -0.58* |
| Maturity |  |  |  | 0.61** | 0.52** | 0.05 | 0.10 | 0.64** | 0.09 | -0.01 |
| Mass |  |  |  |  | 0.53** | 0.38 | -0.19 | 0.64** | -0.19 | 0.32 |
| Longevity |  |  |  |  |  | -0.24 | 0.26 | 0.88** | 0.36 | -0.32 |
| $\pi_S$ | | | | | | | -0.78** | -0.04 | -0.67* | 0.92** |
| $\pi_N/\pi_S$ | | | | | | | | 0.11 | 0.57 | -0.73** |
| Gen. time |  |  |  |  |  |  |  |  | 0.29 | -0.17 |
| $u$ | | | | | | | | | | -0.89** |

Asterisks indicate strength of support (\*\*: $pp > 0.975$ , \*: $pp > 0.95$ )

Although not conclusive, the correlation patterns between *π_S_*, *π_N_ /π_S_* and *dN/dS* are compatible with the hypothesis that the nearly-neutral model is essentially valid for primates. The overall lack of correlation with *LHT*, on the other hand, suggest that there is no clear correlation between effective population size and body size or other related life-history traits in this group. Possibly, the phylogenetic scale might be too small to show sufficient variation in LHT that would be interpretable in terms of variation in *N_e_*. Alternatively, *N_e_* might be driven by other life-history characters (in particular, the mating systems), which may not directly correlate with body size. Of note, in those cases where the estimated correlation of *dN/dS* or *π_N_ /π_S_* with LHTs were in agreement with the predictions of the nearly-neutral theory (Eyre-Walker *et al.*, 2002; Popadin *et al.*, 2007; Nikolaev *et al.*, 2007; Lartillot, 2013; Nabholz *et al.*, 2013; Romiguier *et al.*, 2014; Figuet *et al.*, 2016), the reported correlation strengths were often weak, weaker than the correlations found directly between *π_S_* and *π_N_ /π_S_* and *dN/dS* (Piganeau & Eyre-Walker, 2009; Romiguier *et al.*, 2014; Figuet *et al.*, 2016; Galtier, 2016; Chen *et al.*, 2017; James *et al.*, 2017).

### Teasing apart divergence times, mutation rates and effective population size

The correlation patterns shown by the three molecular quantities *π_S_*, *d_N_ /d_S_* and *π_N_ /π_S_* suggest that *N_e_* plays a non-negligible role in their interspecific variation. However, in its current form, this correlation analysis does not give any quantitative insight about the scaling of *d_N_ /d_S_* and *π_N_ /π_S_* as a function of *N_e_* and, more generally, about the quantitative impact of *N_e_* on the evolution of coding sequences. In order to achieve this, an explicit estimate of the key parameter *N_e_*, and of its variation across species, is first necessary. In this direction, a first simple but fundamental equation relates *π_S_* with *N_e_*:

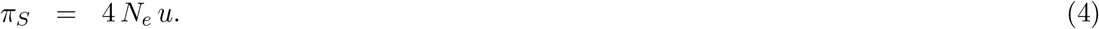

In order to estimate *N_e_* from equation 4, an estimation of *u* is also required. Here, it can be obtained by noting that:

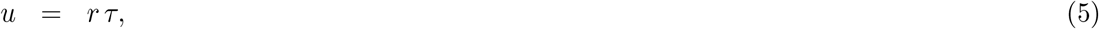

where *r* is the mutation rate per site and per year and *τ* the generation time. Assuming that synonymous mutations are neutral, we can identify the mutation rate with the synonymous substitution rate *dS*, thus leading to:

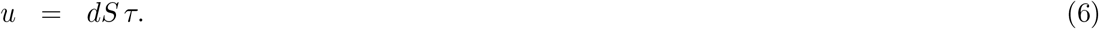

Finally, combining equations 4 and 6 and taking the logarithm gives:

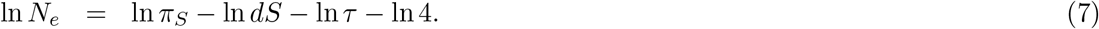

This expression suggests to apply the linear transformation given by equation 7 to the three variables ln *π_S_*, ln *d_S_* and ln *τ*, all of which are jointly reconstructed across the tree by the phenomenological model introduced above, which then gives a global phylogenetic reconstruction of ln *N_e_*. A similar argument, based on equation 6, gives:

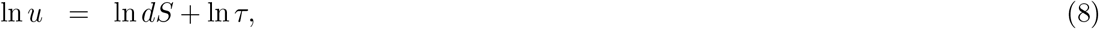

which can be used to obtain a global reconstruction of the mutation rate per generation *u*. Finally, since the two transformations given by equations 7 and 8 are linear, the correlation patterns between ln *N_e_*, ln *u* and the other variables included in the analysis can be recovered by applying elementary matrix algebra to the covariance matrix estimated under the initial parameterization (see Materials and Methods, Ex-post log linear transformation).

The results of this linearly-transformed correlation analysis are gathered in Table 1 (last two columns). First, in accordance with the predictions of the nearly-neutral theory, *π_N_ /π_S_* and *d_N_ /d_S_* show a negative correlation with *N_e_* (*r* = *−*0.73 for *π_N_ /π_S_* and *−*0.58 for *dN/dS*). Second, *u* is inferred to covary negatively with *N_e_* (*r* = *−*0.89), suggesting that species with large *N_e_* tend to have a lower *u* (Lynch *et al.*, 2011). This correlation should be interpreted with caution, however, because these two variables are both estimated partly based on *π_S_* = 4*N_e_u*, such that estimation errors on *π_S_* will systematically affect them in opposite directions. On the other hand, and perhaps more convincingly, *u* is negatively correlated with *π_S_* (*r* = *−*0.67, table 1). Unlike *N_e_* and *u*, *π_S_* and *u* are empirically independent in the present case (i.e. estimated based on different data sources). This latter correlation thus further strengthens the case that *N_e_* and *u* have opposite trends. It also suggests that the variation in *u* is more moderate than the variation in *N_e_*, such that *π_S_* remains in the end positively correlated with *N_e_*.

### Quantitative scaling of *π_N_ /π_S_* and *d_N_ /d_S_* as a function of *N_e_*

Once an explicit estimate of *N_e_* and of its variation is available, the scaling behavior of *π_N_ /π_S_* and *d_N_ /d_S_* as a function of *N_e_* can be quantified. Mathematically, the Brownian process followed by rates and traits, such as given by equation 1, implies the following log-linear relations:

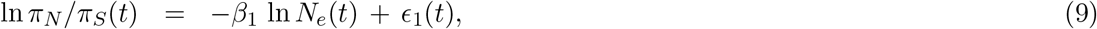

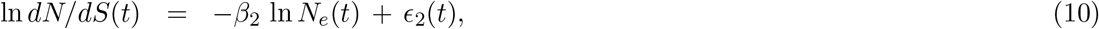

where *ϵ_i_*(*t*), for *i* = 1, 2, are Brownian motions. These last two terms are mathematically equivalent to the residual errors of the linear regression between the independent contrasts of ln *π_N_ /π_S_* and ln *dN/dS* against log *N_e_*. Equivalently:

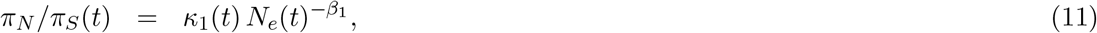

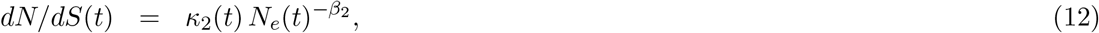

where *κ_i_*(*t*) = *e*^*ϵi*(*t*)^, for *i* = 1, 2. In other words, the slopes of the log-linear relations, *β*_1_ and *β*_2_, are just the scaling coefficients of *π_N_ /π_S_* and *d_N_ /d_S_* as a function of *N_e_*. These two scaling coefficients can be directly obtained based on the covariance matrix estimated above (see methods, Correlations and slopes). Of note, equations 9 to 12 are just a reformulation of the output of the correlation analysis conducted previously. As such, they do not entail any specific hypothesis about the population-genetic relations between *π_N_ /π_S_*, *dN/dS* and *N_e_*. A population-genetic interpretation of these equations is considered in the next subsection.

In the present case, the estimates of *β*_1_ and *β*_2_ are of similar magnitude, with point estimates of 0.17 and 0.10, respectively (Table 2). It is worth comparing these estimates with those that would be obtained if we were using *π_S_* directly as a proxy of *N_e_* (i.e. without correcting for *u*). The slopes of the log-linear scaling of *π_N_ /π_S_* and *dN/dS* as a function of *π_S_* are steeper than those obtained as a function of *N_e_*, with point estimates of 0.29 and 0.13 (Table 2), suggesting that the confounding effects of *u* are not negligible on the evolutionary scale of a mammalian order such as primates and, as such, can substantially distort the scaling relations if not properly taken into account.

**Table 2.** Scaling coefficient of *dN/dS* and *π_N_ /π_S_* as functions of *N_e_* and *π_S_*, compared with estimates of the shape parameter *β* of the distribution of fitness effects.

| method or source | point estimate | credible/confidence interval |
| --- | --- | --- |
| $\pi_N/\pi_S \sim N_e$ | 0.17 | (0.02, 0.3) |
| $dN/dS \sim N_e$ | 0.10 | (-0.02, 0.22) |
| $\pi_N/\pi_S \sim \pi_S$ | 0.29 | (0.12, 0.51) |
| $dN/dS \sim \pi_S$ | 0.13 | (-0.12, 0.51) |
| $\beta$ (mechanistic model) | 0.27 | (0.22, 0.33) |
| Eyre-Walker *et al.* (2006) | 0.23 | (0.19, 0.27) |
| Boyko *et al.* (2008) | 0.18 | (0.16, 0.21) |
| Castellano *et al.* (2019) | 0.16 | (0.13, 0.17) |

### Relation with the shape of the distribution of fitness effects

Mechanistically, the slope of ln *π_N_ /π_S_* and ln *d_N_ /d_S_* as a function of ln *N_e_* can be interpreted in the light of an explicit mathematical model of the nearly-neutral regime. Such mathematical models, which are routinely used in modern Mac-Donald Kreitman tests (Charlesworth & Eyre-Walker, 2008; Eyre-Walker & Keightley, 2009; Halligan *et al.*, 2010; Galtier, 2016), formalize how mutation, selection and drift modulate the detailed patterns of polymorphism and divergence. In turn, these modulations depend on the shape of the distribution of fitness effects (DFE) over non-synonymous mutations (Eyre-Walker & Keightley, 2007). Mathematically, the DFE is often modelled as a gamma distribution. The shape parameter of this distribution (usually denoted as *β*) is classically estimated based on empirical synonymous and non-synonymous site frequency spectra. Typical estimates of the shape parameter are of the order of 0.15 to 0.20 in humans (Boyko *et al.*, 2008; Eyre-Walker *et al.*, 2006), thus suggesting a strongly leptokurtic distribution, with the majority of mutations having either very small or very large fitness effects.

When the shape parameter *β* is small, both *π_N_ /π_S_* and *d_N_ /d_S_* are theoretically predicted to scale as a function of *N_e_* as a power-law, with a scaling exponent equal to *β* (Kimura, 1979; Welch *et al.*, 2008), i.e.:

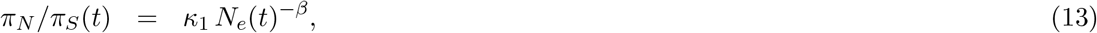

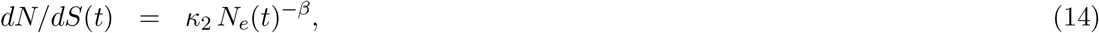

The relation given by equation 13 was used previously for analyzing the impact of the variation in *N_e_* along the genome in *Drosophila* (Castellano *et al.*, 2018). Assuming that (1) the sequences follow a purely nearly-neutral regime, thus without positive selection; (2) the DFE is constant across primate species; (3) variation in *N_e_* is sufficiently slow, compared to the fixation time of non-synonymous substitutions, and (4) short-term *N_e_* and long-term *N_e_* are identical, equations 13 and 14 also predict the interspecific variation in *π_N_ /π_S_* and *dN/dS* as a function of *N_e_*. More specifically, by identifying these two equations with equations 11 and 12, the present argument predicts that the interspecific allometric scaling coefficients *β*_1_ and *β*_2_ estimated above should both be equal to the shape parameter *β* of the DFE. In the present case, the two estimates *β*_1_ and *β*_2_ are indeed congruent with each other, with overlapping credible intervals (Table 2). They are also compatible with previously reported independent estimates of the shape parameter of the DFE, such as obtained from site frequency spectra in Humans and in other great apes (also reported in Table 2).

### A mechanistic nearly-neutral phylogenetic codon model

Since all of the results presented thus far are compatible with a nearly-neutral regime, we decided to construct a mechanistic version of the model directly from first principles. Thus far, a phenomenological approach was adopted, in which the whole set of variables of interest (*dS*, *dN/dS*, *π_S_*, *π_N_ /π_S_* and LHT) were jointly reconstructed along the phylogeny, as a multivariate log-normal Brownian process with 8 degrees of freedom. Only in a second step were *N_e_* and *u* extracted from this multivariate process, using log-linear relations. As a result, *dN/dS*, *π_S_* and *π_N_ /π_S_* are correlated with *N_e_* and *u*, but they are not deterministic functions of these fundamental variables, such as suggested by equations 6, 7, 13 and 14.

In the mechanistic model now introduced, these deterministic relations are enforced. The Brownian process now has 6 degrees of freedom (*π_S_*, *π_N_ /π_S_* and the LHT), corresponding to the variables that are directly observed in extant species. Then, equations 6, 7, 13 and 14 are inverted, so as to express *r* = *dS*, *dN/dS*, *N_e_* and *u* as time-dependent deterministic functions of this Brownian process (see Materials and Methods, Mechanistic model). In these equations, the shape parameter *β*, along with *κ*_1_ and *κ*_2_ in equations 13 and 14, are structural parameters of the model, representing the DFE, itself assumed to be constant across species. These parameters are estimated along with the covariance matrix, divergence times and the realization of the process along the phylogeny. The model is conditioned on the same data (sequence alignment, traits, polymorphism) and using the same fossil calibration scheme as for the phenomenological model.

This mechanistic model was implemented in two alternative versions. A first naive version assumes that genetic divergence takes place exactly at the splitting times defined by the underlying species phylogeny. Identifying genetic divergence with species divergence, however, amounts to ignoring the time it takes for coalescence to occur in the ancestral populations. For that reason, an alternative model was considered, based on the argument that the mean coalescence time in the ancestral population at a given ancestral node of the phylogeny is equal to *δt* = 2*N_e_τ*, where *N_e_* is the effective population size and *τ* the generation time prevailing at or around that node of the species tree. Since *N_e_* and *τ* are both reconstructed along the entire species phylogeny, it is therefore possible to use the local information about the current value of *N_e_* and *τ* to account for the extra amount of divergence *δt* induced by coalescence in the ancestral population when calculating the sequence likelihood (see methods). In the following, these two alternative versions of the mechanistic model are called the ‘naive-phylogenetic’ and ‘mean-coalescent’ versions. Of note, the mean-coalescent version is solely invoked to correct mutation rate and time estimates, not *dN/dS*. In reality, ancestral coalescence is expected to lead to non-trivial patterns of non-synonymous and synonymous divergence (Mugal *et al.*, 2020), which are ignored here.

Fitting the model on the data returns an estimate of the structural parameters of the DFE as well as a dated tree, annotated with the complete history of the variation in mutation rate and in effective population size along its branches. These different aspects of the output of the analysis are now considered in turn.

### Mechanistic estimate of the shape parameter of the DFE

The mechanistic model yields an estimate of *β* (Table 2) that is somewhat higher than the slopes *β*_1_ and *β*_2_ estimated previously with the phenomenological approach, with a posterior median at 0.27 and a credible interval equal to (0.22, 0.33). The credible interval is smaller than those for *β*_1_ and *β*_2_, reflecting the fact that the mechanistic model is more constrained. Our estimate of *β* also appears to be higher than independent estimates obtained from site frequency spectra. Of note, there is some discrepancy among those DFE-based estimates, whose credible intervals do not overlap. However, our estimate is significantly higher than the more recent estimate obtained by jointly analyzing the site frequency spectra across great apes (Castellano *et al.*, 2019), which is probably the most relevant one in the present context. This observation suggests a potential violation of the mechanistic model (see Discussion).

### Estimated mutation rates

The dated phylogeny, together with the reconstructed history of the mutation rate per year *r*, is shown in Figure 1. Here, the mechanistic model works like a standard molecular dating method, using fossil calibration and a relaxed clock model to tease out times and rates from raw synonymous sequence divergence. Generation time, also reconstructed along the tree by the model, is then used to convert the mutation rate per year *r* into a mutation rate per generation *u* (Figure 2). Point estimates (medians) and 95% credible intervals of both *r* and *u* for some extant species of interest and a few key ancestors are shown in Table 3.

**Figure 1.**
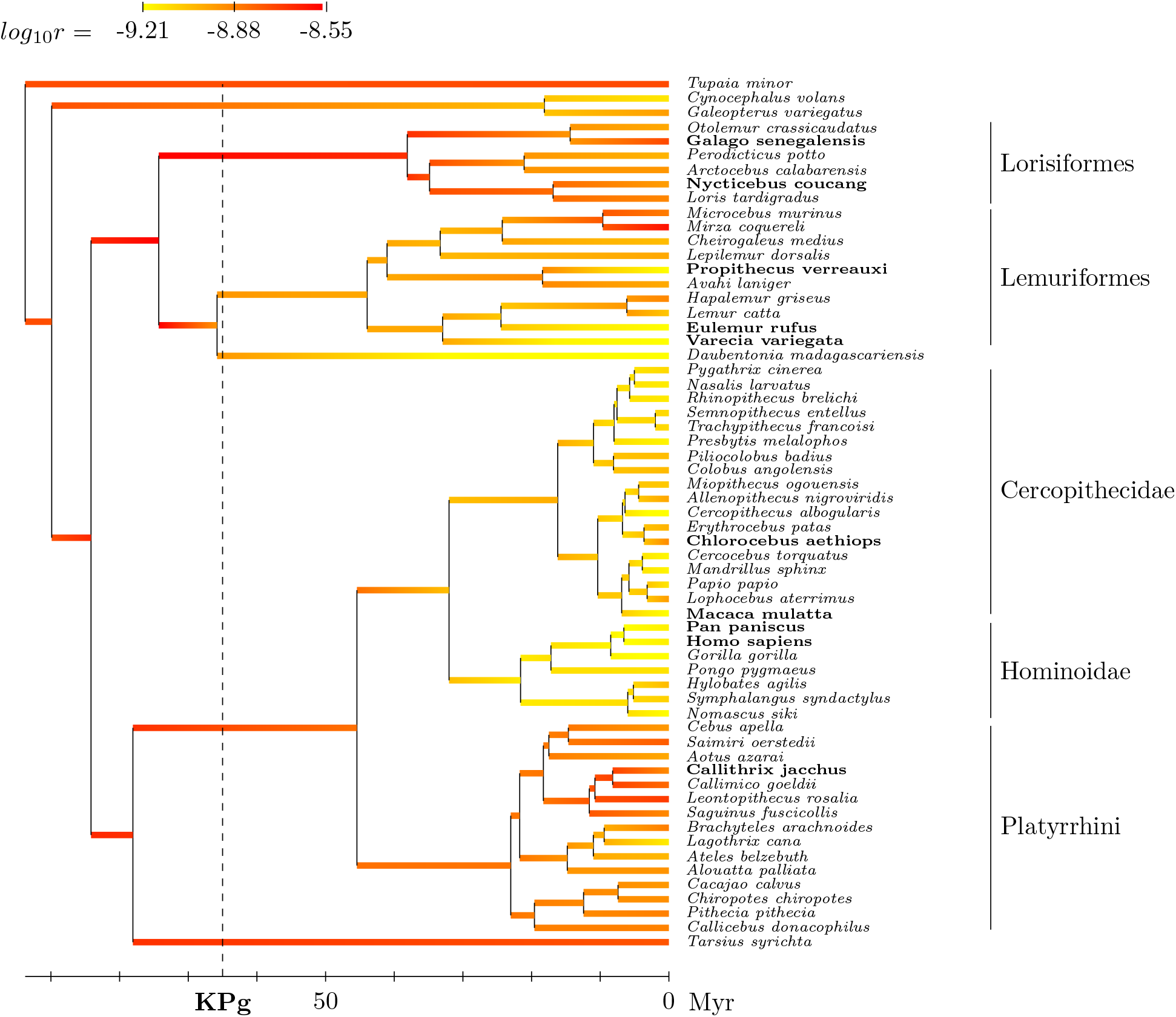
Reconstructed phylogenetic history of the mutation rate per year *r* (posterior median estimate) under the mechanistic model. Species for which transcriptome-wide polymorphism data were used are indicated in bold face.

**Figure 2.**
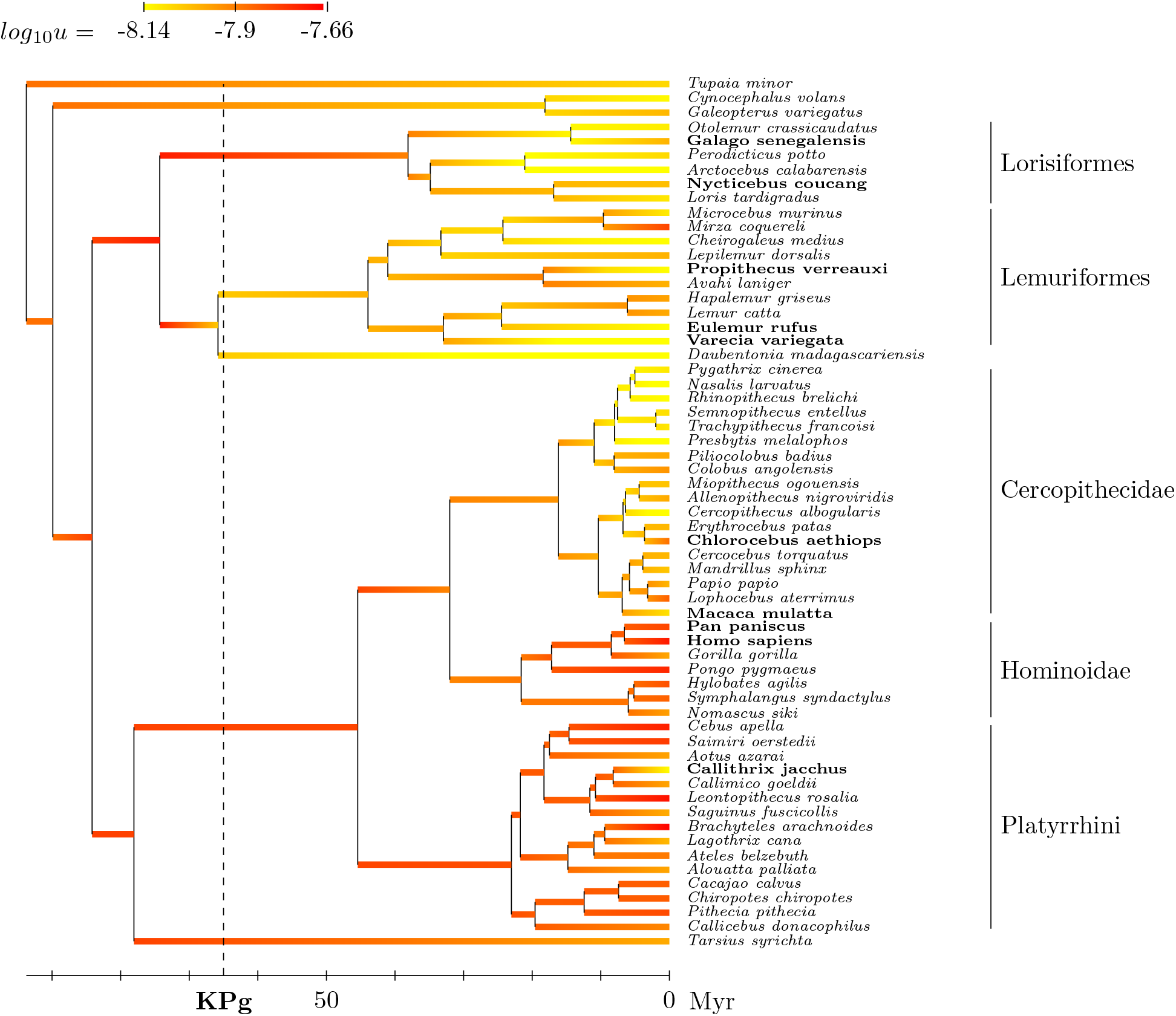
Reconstructed phylogenetic history of the mutation rate per generation *u* (posterior median estimate) under the mechanistic model. Species for which transcriptome-wide polymorphism data were used are indicated in bold face.

**Table 3.** Estimates of mutation rate per year *r* and per generation *u* (posterior median and 95% credible interval), for several extant and ancestral species.

| species | $r$ (per $10^9$ years) | | $u$ (per $10^8$ generation) | | pedigrees<sup>c</sup> |
| --- | --- | --- | --- | --- | --- |
|  | without anc. pol.<sup>a</sup> | with anc. pol.<sup>b</sup> | without anc. pol.<sup>a</sup> | with anc. pol.<sup>b</sup> |  |
| *Homo* | 0.81 ( 0.61, 1.11) | 0.67 ( 0.50, 0.91) | 2.36 ( 1.76, 3.21) | 1.95 ( 1.46, 2.65) | 1.23 - 1.29 |
| *Pan* | 0.79 ( 0.67, 0.93) | 0.67 ( 0.56, 0.80) | 1.91 ( 1.61, 2.23) | 1.62 ( 1.34, 1.92) | 1.26 - 1.48 |
| *Gorilla* | 0.73 ( 0.44, 1.13) | 0.55 ( 0.32, 0.85) | 1.39 ( 0.84, 2.15) | 1.05 ( 0.60, 1.62) | 1.13 |
| *Pongo* | 0.84 ( 0.54, 1.20) | 0.73 ( 0.46, 1.04) | 2.11 ( 1.36, 3.00) | 1.84 ( 1.16, 2.59) | 1.66 |
| *Macaca* | 0.68 ( 0.54, 0.81) | 0.58 ( 0.46, 0.71) | 0.95 ( 0.75, 1.14) | 0.82 ( 0.64, 1.00) | 0.37 |
| *Papio* | 0.90 ( 0.50, 1.59) | 0.71 ( 0.38, 1.30) | 1.23 ( 0.77, 1.94) | 0.98 ( 0.57, 1.59) | 0.55 |
| *Aotus* | 1.09 ( 0.63, 1.77) | 1.02 ( 0.56, 1.74) | 1.16 ( 0.70, 1.78) | 1.08 ( 0.61, 1.70) | 0.81 |
| Catarrhini | 0.87 ( 0.54, 1.45) | 0.91 ( 0.55, 1.52) | 1.24 ( 0.83, 1.79) | 1.23 ( 0.82, 1.85) |  |
| Platyrrhini | 1.48 ( 1.00, 2.15) | 1.42 ( 0.96, 2.09) | 1.57 ( 1.18, 2.15) | 1.53 ( 1.12, 2.06) |  |
| Haplorrhini | 2.00 ( 1.01, 4.07) | 2.09 ( 0.98, 4.42) | 1.44 ( 0.88, 2.38) | 1.60 ( 0.92, 2.84) |  |
| Strepsirrhini | 2.57 ( 1.34, 4.99) | 2.77 ( 1.29, 5.94) | 1.65 ( 1.02, 2.73) | 1.88 ( 1.08, 3.38) |  |
| Primates | 2.07 ( 1.06, 4.24) | 2.20 ( 1.05, 4.79) | 1.45 ( 0.87, 2.47) | 1.62 ( 0.94, 2.93) |  |

<sup>a</sup> naive-phylogenetic method (not accounting for ancestral polymorphism)
<sup>b</sup> mean-coalescent method (accounting for ancestral polymorphism)
<sup>c</sup> from Table 1 of Wu *et al.* (2020)

The naive-phylogenetic model tends to return higher mutation rates, compared to the mean-coalescent approach, in particular for extant taxa and, more generally, for recent ancestors. For instance, in humans, the mutation rate per year is estimated at 0.80 *×* 10^*−*9^ when coalescence in the ancestral population is ignored, versus 0.67 *×* 10^*−*9^ when it is accounted for in the model. This reflects a bias induced by ignoring ancestral coalescence. For instance, the Human-Chimpanzee split is constrained at around 5 to 7 My, but coalescence in the ancestral population can easily go back to 9 My, which, if dated at 7 My, automatically induces an overestimation of the mutation rate along the branch leading to Humans (Besenbacher *et al.*, 2019).

The estimates obtained here under the mean-coalescent model are intermediate, lower than previously reported phylogenetic estimates but still higher than pedigree-based estimates. For instance, typical phylogenetic estimates for the mutation rate in humans are typically of the order of 10^*−*9^ per year, or 3 *×* 10^*−*8^ per generation, whereas pedigree-based estimates are generally about half these values. Concerning other primates, our estimates are also higher than pedigree based estimates previously reported for macaques (Wang *et al.*, 2020) and baboons (Wu *et al.*, 2020). On the other hand, they are congruent for Gorilla, Pongo (Besenbacher *et al.*, 2019) and Aotus (Thomas *et al.*, 2018). In the following, only coalescent-aware estimates are further considered.

Across primates, the mutation rate per year *r* shows a 5-fold variation from 0.6 *×* 10^*−*9^ to 3.0 *×* 10^*−*9^ point mutations per year and per nucleotide site (Figure 1). The rate of mutation is relatively high in the ancestor of primates (*~* 2 *×* 10^*−*9^). On the side of Strepsirrhini, it remains high in Lorisiformes (*~* 2 *×* 10^*−*9^) but is lower in Lemuriformes (*~* 10^*−*9^). Concerning Haplorrhini, the rate undergoes a net slowdown in Catarrhini (*~* 10^*−*9^), further accentuated in apes, which are among the slowest evolving primates (*~* 0.6 *×* 10^*−*9^). These observations are globally in accordance with previous observations, in particular, emphasizing that the slowdown occurring in apes (Steiper *et al.*, 2004) is in fact in the continuity of a broader process of deceleration more generally across catarrhine primates (Perelman *et al.*, 2011).

Compared to the mutation rate per year, the mutation rate per generation *u* varies over a more moderate range, showing a 3-fold variation across primates. Moderately high ancestrally (1.5 *×* 10^*−*8^), it shows a convergent decrease in Lorisiformes, Lemuriformes, and Cercopithecidae (reaching below 10^*−*8^ in several species in these three clades), but otherwise, remains more in the range of 1.5 to 2 *×* 10^*−*8^. At first sight, the mutation rates per year *r* and per generation *u* tend to show opposite patterns: slow-evolving lineages, with a low mutation rate per year, tend to have a higher mutation rate per generation (high *u*). This is particularly apparent for apes (low *r*, high *u*) or for Lorisiformes (high *r*, low *u*). However, there are some exceptions, of lineages that have both a high *r* and a high *u*, most notably Platyrrhini (new world monkeys). More generally, the correlation analysis (table 1) suggests that *u* and *r* are positively, not negatively, correlated on average across primates.

### Phylogenetic Reconstruction of *N_e_*

The marginal reconstruction of *N_e_* along the phylogeny of primates returned by the mechanistic model is shown in Figure 3 (Supplementary Figure 1 for the reconstruction returned by the phenomenological model). More detailed information, with credible intervals, is given in Table 4 for several species of interest and key ancestors along the phylogeny.

**Figure 3.**
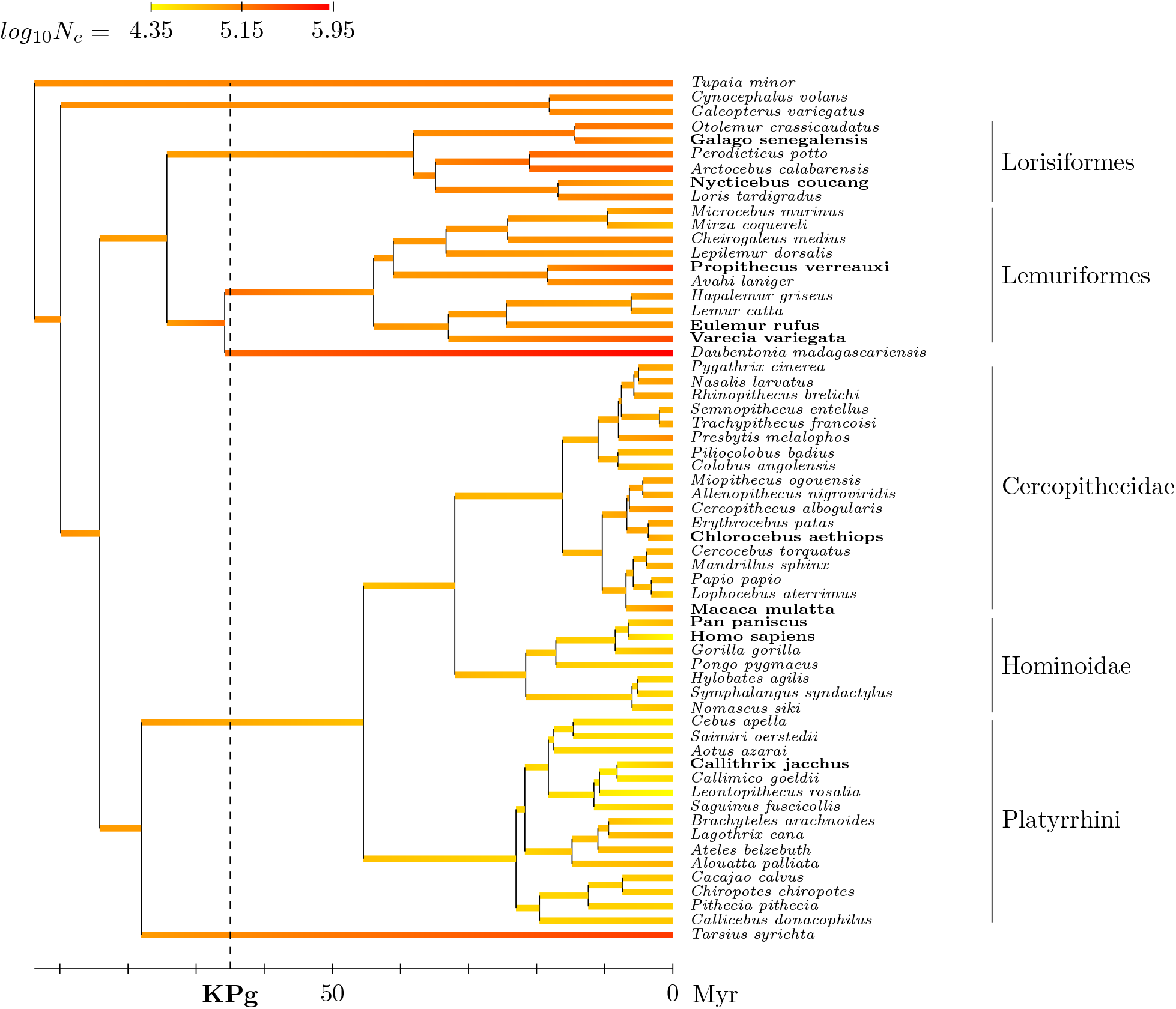
Reconstructed phylogenetic history of *N_e_* (posterior median estimate) under the mechanistic model. Species for which transcriptome-wide polymorphism data were used are indicated in bold face.

**Table 4.** Estimates of effective population size (*×*10^*−*3^, posterior median and 95% credible interval) for several extant taxa and ancestors.

| species | mechanistic | mech. w/o anc. pol. | phenomenological | uncoupled | coal.<sup>a</sup> | hmmcoal<sup>b</sup> |
| --- | --- | --- | --- | --- | --- | --- |
| *Homo* | 23 ( 17, 30) | 19 ( 14, 25) | 19 ( 12, 36) | 20 ( 13, 32) | (13, 16) | 8 |
| *Pan* | 64 ( 54, 78) | 54 ( 46, 64) | 67 ( 45, 101) | 60 ( 40, 94) | (31, 62) | 30 |
| *Gorilla* | 64 ( 25, 170) | 67 ( 27, 159) | 110 ( 37, 354) | 47 ( 20, 108) | (28, 57) | 21 |
| *Pongo* | 42 ( 15, 109) | 38 ( 15, 89) | 52 ( 20, 131) | 32 ( 10, 103) | (42, 85) | 19 |
| *Homo-Pan* | 44 ( 28, 69) | 43 ( 28, 66) | 49 ( 29, 87) | 39 ( 24, 64) | (10, 47) | 50 |
| *Homo-Gorilla* | 45 ( 26, 73) | 46 ( 28, 74) | 54 ( 31, 101) | 41 ( 24, 70) | (27, 61) | 47 |
| Hominidae | 48 ( 24, 91) | 44 ( 24, 78) | 52 ( 26, 104) | 45 ( 21, 97) |  |  |
| Hominoidea | 56 ( 28, 111) | 52 ( 28, 98) | 63 ( 31, 139) | 53 ( 24, 121) |  |  |
| Cercopithecidae | 64 ( 34, 114) | 70 ( 41, 118) | 81 ( 43, 156) | 72 ( 36, 147) |  |  |
| Catarrhini | 67 ( 33, 137) | 68 ( 36, 128) | 70 ( 34, 157) | 57 ( 26, 128) |  |  |
| Platyrrhini | 42 ( 22, 79) | 37 ( 20, 67) | 32 ( 16, 60) | 32 ( 14, 79) |  |  |
| Simiiformes | 56 ( 26, 130) | 53 ( 26, 110) | 43 ( 18, 97) | 39 ( 15, 101) |  |  |
| *Tarsius* | 392 ( 80, 2220) | 285 ( 74, 1477) | 89 ( 18, 391) | 49 ( 4, 568) |  |  |
| Haplorrhini | 96 ( 40, 240) | 88 ( 39, 199) | 46 ( 15, 131) | 45 ( 14, 149) |  |  |
| Lorisiformes | 117 ( 55, 253) | 115 ( 58, 244) | 53 ( 16, 135) | 53 ( 21, 135) |  |  |
| Lemuriformes | 102 ( 47, 218) | 97 ( 48, 197) | 118 ( 48, 285) | 152 ( 66, 377) |  |  |
| *Daubentonia* | 863 ( 162, 6174) | 893 ( 183, 5296) | 739 ( 142, 5583) | 335 ( 43, 2796) |  |  |
| Strepsirrhini | 93 ( 37, 253) | 84 ( 36, 200) | 39 ( 12, 116) | 45 ( 16, 132) |  |  |
| Primates | 97 ( 40, 251) | 88 ( 39, 200) | 47 ( 14, 132) | 46 ( 15, 151) |  |  |

<sup>a</sup> from Prado-Martinez et al, 2013, table 1, for extant hominids, and from Rannala and Yang, 2003 for ancestral species; <sup>b</sup> from Prado-Martinez et al, 2013, figure 2

The *N_e_* estimates for the four extant hominids (*Homo, Pan, Gorilla and Pongo*) are globally congruent with independent estimates based on other coalescent-based approaches (Prado-Martinez *et al.*, 2013). In particular, for Humans, *N_e_* is estimated to be between 13 000 and 24 000. For other hominids, they tend to be somewhat higher than coalescent-based estimates. Concerning the successive last common ancestors along the hominid subtree, our estimation is also consistent with the independent-locus multi-species coalescent (Rannala & Yang, 2003).

The history of *N_e_* shows a clear large-scale structure over the primate phylogeny. Starting with a point estimate at around 100 000 in the last common ancestor of primates, *N_e_* then goes down in Haplorrhini, stabilizing at around 65 000 in Cercopithecidae (old-world monkeys), 50 000 in Hominoidea (going further down more specifically in Humans), and 40 000 in Platyrrhini (new-world monkeys). Conversely, in Strepsirrhini, *N_e_* tends to show higher values, staying at 100 000 in Lemuriformes and going up to 160 000 to 200 000 in Lorisiformes. Finally, rather large effective sizes are estimated for the two isolated species *Tarsius* and *Daubentonia* – although the credible intervals are very large (Table 4). These estimates may not be so reliable, owing to the very long branches leading to these two species.

The reconstruction of *N_e_* shown in Figure 3 mirrors the patterns of *dN/dS* estimated over the tree (Supplementary Figure 2). This is partially expected given the fact that the model relies on the scaling relation between *dN/dS* and *N_e_* given by equation 14. However, the model integrates other sources of information, in particular, from *π_S_* and *π_N_ /π_S_*. In order to get some insight about how each of these variables informs the reconstructed history of *N_e_*, two additional models were considered.

First, an alternative version of the phenomenological model was used (Supplementary Figure 3), in which all time-dependent variables are assumed to evolve independently. Under this uncoupled model, *N_e_* is not informed by *dN/dS* or *π_N_ /π_S_*, but only by *π_S_* = 4*N_e_u* and by the estimates of *u* implied by the relaxed clock and data about generation times. Compared to the mechanistic version just presented, this uncoupled model gives lower *N_e_* estimates most notably for the last common ancestor of primates (30 000, versus 100 000 under the mechanistic model), but also for the two species *Daubentonia* and *Tarsius* (which have a low *dN/dS*). Conversely, it returns higher *N_e_* estimates in Lemuriformes (which have a high *dN/dS* compared to Lorisidae).

Interestingly, under the uncoupled model, there is a substantial uncertainty about the estimation of *N_e_* across the tree: the 95% credible intervals span one order of magnitude on average. This uncertainty is reduced under the reconstructions relying on the additional information contributed by *dN/dS* and *π_N_ /π_S_*, quantitatively, by 30% under the phenomenological, and by 50% under the mechanistic covariant models (Table 4). In the end, there is thus on average a factor 5 between the lower and the upper bound of the 95% credible intervals on *N_e_* estimates under the most constrained (mechanistic) model. Concerning the deep branches of the tree, most of this reduction in uncertainty is primarily contributed by *dN/dS* – which thus gives an idea of how much information is extracted from multiple sequence alignments by these models about ancient population genetic regimes.

Second, as mentioned above, the mechanistic model infers a value of 0.27 for the shape parameter *β*, which is higher than recent SFS-based estimates, of the order of 0.16. Fixing *β* = 0.16 in the mechanistic model returns a reconstruction of *N_e_* over the phylogeny (Supplementary Figure 4) qualitatively similar to that returned by the unconstrained version of the model (Figure 3), although covering a broader range (about 100-fold) than under either the mechanistic or the uncoupled models (about 10- to 30-fold, Figure 3 and Supplementary Figure 3). This illustrates the key role of *β* in calibrating the transfer of information from *dN/dS* to *N_e_*. According to the scaling relation given by equation 14, when *β* is small, a large variation in *N_e_* results in a small shift in *dN/dS*. Thus, conversely, with a smaller *β*, the empirically observed variation in *dN/dS* over the tree implies a broader range of *N_e_* variation across primates. In the present case, this argument also suggests an explanation of why the small value of *β* inferred by SFS is rejected by the mechanistic model – because the variation in *N_e_* implied by the history of *dN/dS* under such a small value of *β* would exceed the one implied by the range of *π_S_* observed in extant species.

Of note, the *N_e_* estimates are lower under the naive-phylogenetic (Supplementary Figure 5) than under the mean-coalescent approach (Figure 3). This difference between the two models is due to the fact that, given *π_S_*, any bias in the estimation of *u* has to be compensated for by an opposite bias of the same magnitude in the estimation of *N_e_*. These discrepancies are relatively minor, however, and the global patterns of the history of *N_e_* along the tree are very similar in both cases.

Finally, the phylogenetic history of the mutation rate per generation *u* mirrors that of *N_e_*, such that species with smaller *N_e_* tend to have a higher value for *u* (compare Figures 1 and 2). These opposite patterns of variation, combined with the fact that *N_e_* shows a greater amplitude in its variation across primates compared to u, results in *π_S_* being mostly driven by *N_e_*, although with a partial dampening of its overall variation. This joint pattern for *N_e_* and *u* explains why the regression slopes of ln *π_N_ /π_S_* and ln *dN/dS* against *π_S_* are steeper than those against *N_e_* (Table 2).

## Discussion

### A Bayesian integrative framework for comparative population genomics

The question of the role of *N_e_* in the evolution of coding sequences has motivated much work over the years. One main problem that has attracted particular attention is to understand to what extent *N_e_* modulates the ratio of non-synonymous over synonymous polymorphism (*π_N_ /π_S_*) or divergence (*d_N_ /d_S_*). Often, *π_S_* has been used as a proxy for *N_e_*. However, *π_S_* also depends on *u*, the mutation rate per generation, which differs between species.

In this context, the main contribution of the present work is to propose a Bayesian integrative phylogenetic framework for conducting such comparative analyses in a way that allows for direct quantitative estimation of *N_e_* and of its impact on molecular evolution across a clade of interest. Relying on an integrated relaxed clock model to calibrate mutation rates, the program leverages an estimate of *N_e_* based on *π_S_*, correcting for *u*. Simultaneously, it conducts a regression analysis, returning an estimate of the scaling exponent of molecular quantities such as *dN/dS* and *π_N_ /π_S_*, but also potentially other variables or quantitative traits, directly as a function of *N_e_*. As a byproduct, the approach also returns a global reconstruction of the history of effective population size and mutation rate across the phylogeny.

As can be seen from Table 2, correcting for variation in mutation rate between species (for *u*), as opposed to regressing directly against *π_S_*, does have an impact on the estimated scaling relations. In the present case, the slopes as a function of *π_S_* tend to be steeper than as a function of *N_e_*, a pattern that is more generally expected if species with large *N_e_* also tend to have lower mutation rates per generation, such as previously suggested (Lynch *et al.*, 2011) and confirmed by our correlation analysis (Table 1). The approach introduced here should therefore represent a useful methodological contribution in the context of the current discussions on the role played by those scaling coefficients in molecular evolution (James *et al.*, 2017; Castellano *et al.*, 2018, 2019; Galtier & Rousselle, 2020).

### Estimating mutation rates

Conceptually, our approach for extracting *N_e_* is merely a reformulation, in a Bayesian integrative framework, of the classical idea of estimating *N_e_* from *π_S_* by factoring out the mutation rate *u*, itself estimated based on a molecular clock argument. The integrative approach presents several advantages, however. First, it imposes the same assumptions about the molecular clock, relying on the same sequence data and the same global set of fossil constraints, uniformly for all species included in the analysis. Second, it automatically propagates the uncertainty about estimates of *u*, which themselves incorporate the uncertainty about divergence times, onto the credible intervals eventually reported for *N_e_* or for the slopes of the regressions. Third, these slopes are also automatically corrected for phylogenetic non-independence. Finally, on purely practical grounds, the application of the method is straightforward, just requiring as its input a multiple sequence alignment for the clade of interest, estimates of *π_S_* and *π_N_ /π_S_* for some or all of the extant species and fossil calibrations.

An alternative to the phylogenetic estimation of *u* is to rely on high-throughput sequencing of pedigrees. As of yet, such estimates are available only for 7 primates (Chintalapati & Moorjani, 2020), but this is likely to change in the future. For these 7 species, the phylogenetic estimates obtained here are higher than pedigree-based estimates, thus in line with previous observations. The reasons for this discrepancy are not yet well-understood (Scally & Durbin, 2012; Ségurel *et al.*, 2014; Chintalapati & Moorjani, 2020). Interestingly, accounting for coalescence in the ancestral populations contributes a lot to making phylogenetic estimates closer to those obtained in the same species by sequencing pedigrees, although not entirely.

In principle, pedigree-based estimates of *u* could be included in the framework presented here, as additional constraints at the tips of the phylogeny to inform the reconstruction of *N_e_*. However, given the still unexplained mismatch between pedigrees and phylogenies, it is perhaps more meaningful to compare them after the fact, as was done here (Table 3), and then further investigate the various entry points in the model, at the level of the relaxed clock, the prior on divergence times, the fossil constraints, that could be responsible for this discrepancy.

### Mechanistic models of coding sequence evolution

Our phylogenetic approach was implemented in two alternative versions, using either a phenomenological or a mechanistic modeling strategy. The phenomenological model implements the idea of conducting comparative regression analyses directly against *N_e_*, such as discussed above. In itself, this approach is agnostic about the underlying selective regimes over proteins and, in particular, is not inherently committed to a nearly-neutral interpretation.

The mechanistic model, on the other hand, makes more aggressive assumptions about the underlying selective regime. It is fundamentally a Bayesian phylogenetic implementation of the nearly-neutral theory. Accordingly, it assumes that protein-coding sequences are exclusively under purifying selection. Another key assumption of the model, not necessarily implied by the nearly-neutral theory, is that the DFE is constant across species. These assumptions give more constraint to the analysis and return more focussed estimates. However, the estimate of the shape parameter of the DFE obtained under this model turns out to be significantly higher than some of the estimates based on SFS obtained in Humans or in great apes (Table 2), suggesting that one of these two assumptions might not be strictly valid.

The question of whether the DFE is constant across species has recently motivated both methodological work for jointly analyzing the site frequency spectra of multiple species (Tataru & Bataillon, 2019) and empirical investigations (Castellano *et al.*, 2019; Galtier & Rousselle, 2020). These empirical analyses suggest that the shape parameter is rather stable across great apes (Castellano *et al.*, 2019) and more broadly across primates Galtier & Rousselle (2020). The mean of the distribution, on the other hand (usually denoted 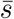), was found to be potentially variable across great apes (Castellano *et al.*, 2019), such that species with larger *N_e_* values also tend to have more strongly deleterious non-synonymous mutations. The rather steep variation of the population scaled mean 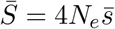 as a function of *π_S_* observed across metazoans Galtier & Rousselle (2020) might also be interpreted as reflecting an underlying positive covariation of 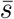 with *N_e_*. Importantly, since *dN/dS* and *π_N_ /π_S_* scale as 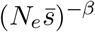, this specific pattern of covariation between the unscaled mean of the DFE and *N_e_* predicts that the regression slopes should be steeper than *β*. This could explain the high estimate of *β* obtained here under the mechanistic model. Of note, the scaling of *π_N_ /π_S_* and *dN/dS* with respect to *N_e_* returned by the phenomenological model are compatible with SFS-based estimates, but this might just be a consequence of the rather large credible intervals obtained in their case.

To further investigate this question, conducting a broader analysis with polymorphism data obtained for a larger number of primate species – and using the same set of coding genes for estimating *π_S_* and *π_N_ /π_S_*, on one hand, and *dS* and *dN/dS*, on the other hand – would certainly be an important direction to pursue, as it would consolidate the results presented here, which are still preliminary in many respects. In particular, it would yield more precise estimates of the scaling coefficients under the phenomenological model. If this confirms the discrepancy between the interspecific scaling coefficients and SFS-based estimates, then, the assumption of a constant DFE under the mechanistic model could be relaxed, although this would then require incorporating information, not just about mean diversity (*π_S_* and *π_N_ /π_S_*), but also about site frequency spectra in extant species, in order to constrain the estimation.

Concerning the other assumption of the mechanistic model, of an exclusively purifying selection regime, the *dN/dS* ratio may in fact contain a fraction of adaptive substitutions, susceptible to distort the relation between *dN/dS* and *N_e_*. The relative importance of adaptive versus nearly-neutral substitutions in primates is still debated (Eyre-Walker & Keightley, 2009; Galtier, 2016; Zhen *et al.*, 2021). In any case, adaptive substitutions might certainly represent an important issue when applying the method to other phylogenetic groups. Here also, the model could be further elaborated, by explicitly including an adaptive component to the total *dN/dS*. Quite interestingly, the resulting model could then be seen as an integrative multi-species version of the Mac-Donald Kreitman test, returning an estimate of the history of the adaptive substitution rate over the phylogeny – which could then be compared with independent estimates based on pairs of sister species (Charlesworth & Eyre-Walker, 2008; Eyre-Walker & Keightley, 2009; Halligan *et al.*, 2010; Galtier, 2016).

Finally, another potential issue, which concerns both the mechanistic and the phenomenological model, is that short-term *N_e_* (such as reflected by *π_S_*) may be strongly dependent on recent demographic events (Charlesworth, 2009) and may thus not be identical with long-term *N_e_* (such as reflected by *d_N_ /d_S_*). This might be one of the reasons why *d_N_ /d_S_* shows a weaker correlation with *π_S_* than *π_N_ /π_S_*. A possible improvement of our model in this direction would consist in allowing for an additional level of variability at the leaves, representing the mismatch between long- and short-term *N_e_*. Other sources of variance in extant diversity estimates could also be modeled, in particular, the additional stochasticity contributed by the random genealogy or by the low counts of SNPs. These last two points are probably a minor issue for nuclear exome-wide polymorphism data, such as explored here. In contrast, they could be quite relevant in the case of the small and non-recombining mitochondrial genome, for which the question of the inter-specific scaling behavior of *dN/dS* and *π_N_ /π_S_* as a function of *N_e_* is also of interest (James *et al.*, 2017).

### Evolution of mutation rates, *N_e_* and life history across primates

The global phylogenetic history of *N_e_* (Figure 3) and mutation rates (Figures 1 and 2) obtained here offers interesting insights into the macro-evolutionary trends in primates, making connections between life-history and molecular evolution. Previous analyses have repeatedly pointed out a slowdown of the molecular clock in apes (Steiper *et al.*, 2004), more broadly in catarrhine primates (Perelman *et al.*, 2011), or even more globally throughout the evolutionary history of the entire order (Steiper & Seiffert, 2012), suggesting a trend towards increasing body size and longer generation times in this group. Our reconstruction confirms this global picture, adding another feature, in the form of a global decrease in effective population size, although more specifically in simians (Figure 3). A global picture only in terms of evolutionary trends along a small-versus-large body size axis, however, would be an oversimplification. In particular, *dS* and *dN/dS* appear to respond differently to LHT, *dS* being negatively correlated with body size (Table 1) as previously reported (Steiper & Seiffert, 2012), whereas *dN/dS* correlates only with longevity but not with body size, a pattern also observed across mammals (Nikolaev *et al.*, 2007; Lartillot, 2013).

The trend in decreasing *N_e_* observed here is primarily driven by the underlying variation in *dN/dS*. As such, it provides another illustration of the more general result that molecular evolutionary patterns inferred from genetic sequences using phylogenetic methods can be informative about life-history evolution (Lartillot & Delsuc, 2012; Romiguier *et al.*, 2013; Figuet *et al.*, 2014; Wu *et al.*, 2017). Compared to previous work, however, an important new contribution of the present work is a quantitative reconstruction, over the phylogeny, directly in terms of the canonical parameters of population genetics, the mutation rate *u* and the effective population size *N_e_*. Such broad-scale reconstructions, as opposed to focussed estimates in isolated extant species, are potentially useful in several respects. First, they provide a basis for further testing some of the key ideas about the role of mutation rate or genetic drift in genome evolution (Lynch *et al.*, 2011; Lefébure *et al.*, 2017). Second, the integrative framework could be augmented with trait-dependent diversification models (Fitzjohn, 2010), so as to examine the role of *N_e_* or *u* in speciation and extinction patterns.

## Materials & Methods

### Coding sequence data, phylogenetic tree and fossil calibration

The coding sequences were taken from Perelman *et al.* (2011) and modified. It consists in a modified subset, codon compliant, based on 54 nuclear autosomal genes in 61 species of primates, and of a total length 15.9 kb. We used the tree topology published by Perelman et al. (itself based on a maximum likelihood analysis), as well as the eight fossil calibrations that were used in this previous study to estimate divergence times. These calibrations were encoded as hard constraints on the molecular dating analysis.

### Life History Traits

We used four life history traits (LHT) in this study. Adult body mass (as a proxy for body mass, 16 missing values), maximum recorded lifespan (ML, as a proxy for longevity, 19 missing values) and female age of sexual maturity (ASM, 26 missing values) were obtained from the An Age database (de Magalhaes & Costa, 2009). Estimates about generation time were calculated from maximum longevity and age at maturity following a method detailed by UICN (Pacifici *et al.*, 2013):

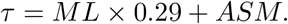

In the case of great apes, and most notably for Humans, these estimates appear to be too high (48.5 years for *Homo*, 24.7 for *Pan*, 23.8 for *Gorilla* and 24.1 for *Pongo*). For these four species, direct estimates of the generation time (29, 24, 19 and 25 years, respectively) were taken from Besenbacher *et al.* (2019).

### Estimation of Polymorphism (*π_S_* and *π_N_ /π_S_*)

The estimates of the synonymous nucleotide diversity *π_S_* and the ratio of non-synonymous over synonymous diversity *π_N_ /π_S_* and *π_S_* of 10 primate species were calculated on the sequence data from Perry *et al.* (2012), such as re-analysed by Romiguier *et al.* (2014) and Figuet *et al.* (2016). We matched these polymorphism data for the three species *Pan troglodytes*, *Propithecus vereauxi coquereli* and *Eulemur mongoz*, to *Pan paniscus*, *Propithecus verreauxi* and *Eulemur rufus*, respectively, from the Perelman et al multiple sequence alignment.

A first series of analyses were conducted using the estimates of *π_N_ /π_S_* and *π_S_* reported by Romiguier *et al.* (2014). Alternatively, and in order to avoid the artifactual correlations induced between *π_S_* and *π_N_ /π_S_* by shared data sampling error, the method of Romiguier *et al.* (2014) was adapted so as to estimate *π_S_* and *π_N_ /π_S_* on different subset of the sites, using the hypergeometric method (James *et al.*, 2017). Specifically, starting from the original fasta file containing the 8 variants for each contig of a given species, coding sites were randomly partitioned into two subsets, with equal probability independently for each site, and the *π_S_* and *π_N_ /π_S_* statistics were computed on each subset using the dNdSpiNpiS software program (available at https://kimura.univ-montp2.fr/calcul/softwares.html), with the same options as those used in Romiguier *et al.* (2014). Finally, the *π_S_* estimate from the first half was combined with the *π_N_ /π_S_* estimate obtained on the second half. The results presented in the main document are all based on this hypergeometric method. They are essentially identical to those obtained using the original polymorphism estimates (Supplementary Table 2).

## Models

### The phenomenological model

The phenomenological model is essentially the one introduced in Lartillot & Poujol (2011), with some minor modifications. The exact structure of the model is given in the manual of Coevol, version 1.5b.

### Correlations and slopes

Given a covariance matrix Σ describing the correlation structure of a Brownian process *X*, the strength of the correlation coefficient between two entries of *X*, *k* and *l*, is given by:

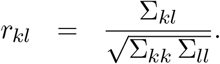

The slope of the regression of trait *l* agains trait *k* is given by:

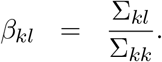

In both cases, the equation is applied successively on each point obtained from the posterior distribution by MCMC, yielding a collection of values, for either the correlation coefficient or the slope, from which a median point estimate and a credible interval are then computed.

### Ex-post log-linear transformation of the correlation analysis

In the following, the multivariate process *X* is specified such that the entries are in the same order as in table 1 (*dS*, *dN/dS*, maturity, mass, longevity, *π_S_*, *π_N_ /π_S_* and generation time *τ*). Based on this vector *X*, an extended vector *Y* can be defined, as:

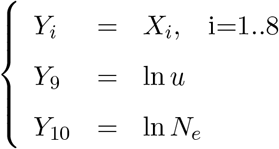

Since ln *u* and ln *N_e_* are given by the following log-linear relations:

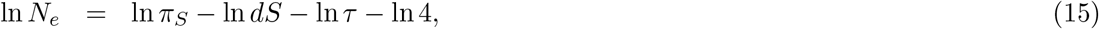

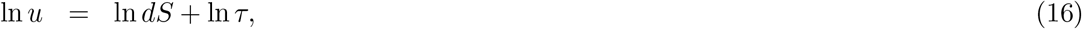

*Y* can be expressed as *Y* = *AX* + *K*. Here, *K* is a vector of constants (depending on the time scale), and *A* is an 8 *×* 10 matrix:

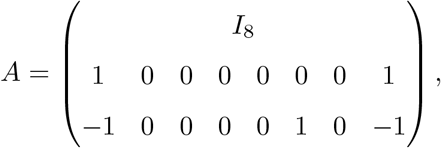

where *I*_8_ is the 8 *×* 8 identity matrix. Since *X* is Brownian, parameterized by a covariance matrix Σ_*X*_, *Y* is also Brownian, with (degenerate) covariance structure given by Σ_*Y*_ = *A ×* Σ_*X*_ × *A′*. In practice, we added a new method in Coevol to read the output and apply the linear transformation (from *X* and Σ_*X*_ to *Y* and Σ_*Y*_) on each sample from the posterior distribution. This gives a reconstruction of *N_e_* (posterior median, credible intervals) and of the correlation matrix Σ_*Y*_.

### Mechanistic Nearly-Neutral Model

This alternative model uses the original Coevol framework but introduces additional constraints, such that some of the parameters are deduced through deterministic relations implying other Brownian dependent parameters. Specifically, the Brownian free variables are now:

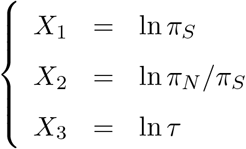

Then, using equations 6, 7, 11 12, the other variables of interest can be expressed as deterministic functions:

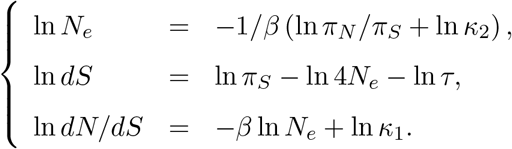

This model has three structural free parameters, *β*, *κ*_1_ and *κ*_2_, which were each endowed with a normal prior, of mean 0 and variance 1.

The naive-phylogenetic version of the model uses the default approach used in Coevol, i.e. assumes a single dated tree *T*, with branch lengths measured in time. The Brownian multivariate process *X* runs along this tree, splitting into two independent processes whenever a speciation node is encountered along the phylogeny. Conditional on *T* and *X*, the sequences then evolve along that same tree *T*, using a codon-model in which *dS* and *dN/dS* are modulated across branches, such as implied by *X* (Lartillot & Poujol, 2011). For the mean-coalescent version of the model, on the other hand, conditional on *T* and *X*, the sequence evolutionary process is assumed to run on a dated tree *T′* different from *T*. Compared to *T*, the node ages of *T′* are all shifted further back into the past by an amount *δt* = *Max*(2*N_e_τ, δt*_0_), where *N_e_* and *τ* are the instant values of effective population size and generation time implied by the process *X* at the corresponding node in the original tree *T*, and *δt*_0_ is the difference between the age of the focal node on *T* and the age of its oldest daughter node in *T′*. This additional constraint is meant to ensure that a node should always be older than its daughter nodes in *T′*. Owing to the variation in *N_e_* and *τ* between successive nodes, this constraint may not be automatically realized, in particular in regions of the tree in which successive splitting times are within coalescence time from each other – precisely those splits that are potentially under a regime of incomplete lineage sorting. In practice, this problem is rarely encountered (less than one node of the whole tree on average under the posterior distribution).

### Uncoupled Model

The *uncoupled* model, already implemented in Coevol, is similar to the phenomenological version of the model, except that the variables of interest (*dS*, *dN/dS*, *π_S_*, *π_N_ /π_S_* and *τ*) are modelled as independent Brownian processes along the tree. Equivalently, we use a multivariate Brownian model with a diagonal covariance matrix (see Lartillot and Poujol, 2011, for details).

### Markov Chain Monte-Carlo (MCMC) and post-analysis

Two independent chains were run under each model configuration. Convergence of the chains was first checked visually (a burnin of approximately 1000 points, out of 6000, was taken for the phenomenological model, and of 100 out of 3100 for the mechanistic model) and quantified using Coevol’s program Tracecomp (effective sample size greater than 500 and maximum discrepancy smaller than 0.10 across all pairs of runs and across all statistics, except for the age of the root of the tree, which typically has a lower effective sample size). We used the posterior median as the point estimate. The statistical support for correlations is assessed in terms of the posterior probability of a positive or a negative correlation. The slope is estimated for each covariance matrix sampled from the distribution, which then gives a sample from the marginal posterior distribution over the slope.

### Software and data availability

Coevol (Lartillot & Poujol, 2011) is an open source program available on github: https://github.com/bayesiancook/coevol. All models and data used here, along with scripts to re-run the entire analysis, are accessible from this repo. Data and scripts for computing *π_S_* and *π_N_*/*π_S_* are available at https://github.com/bayesiancook/polyprim

## Authors contribution

The modifications of Coevol and the new models presented in this study are primarily attributable to MB, who also gathered and formatted the data and conducted all analyses, in the context of an internship (master Biosciences of École Normale Supérieure de Lyon). NL contributed additional analyses and further elaborations for the revised version. MB and NL both contributed to the writing of the manuscript.

## Acknowledgements

We wish to thank Emeric Figuet, Jonathan Romiguier and Nicolas Galtier for sharing polymorphism data (PopPhyl project) and for their help in re-running the scripts for analysing them, Emmanuel Douzery and Frederic Delsuc for editing the multiple sequence alignment of Perelman et al to make it codon-compliant, and Nicolas Galtier, Laurent Duret and Thibault Latrille for their input on this work and their comments on the manuscript.

