## Supplemental Tables and Figures for "Reconstructing the history of variation in effective population size along phylogenies"

June, 5, 2021

**Running head:** A phylogenetic history of $N_e$

Table S1. Correlation coefficients between $dS$, $dN/dS$, $\pi_S$, $\pi_N/\pi_S$, life-history traits and $N_e$ under a uniform prior over divergence times.

| | $dS$ | $dN/dS$ | Mat. | Mass | Long. | $\pi_S$ | $\pi_N/\pi_S$ | Gen. | $u$ | $N_e$ |
|---|---|---|---|---|---|---|---|---|---|---|
| $dS$ | | 0.27 | -0.33 | -0.57** | -0.20 | -0.63* | 0.51 | -0.38 | 0.80** | -0.78** |
| $dN/dS$ | | | 0.16 | 0.10 | 0.50* | -0.49 | 0.42 | 0.43 | 0.57* | -0.57* |
| Maturity | | | | 0.61** | 0.53** | 0.04 | 0.10 | 0.65** | 0.06 | -0.01 |
| Mass | | | | | 0.52** | 0.35 | -0.18 | 0.64** | -0.19 | 0.30 |
| Longevity | | | | | | -0.23 | 0.25 | 0.87** | 0.33 | -0.30 |
| $\pi_S$ | | | | | | | -0.79** | -0.02 | -0.69* | 0.91** |
| $\pi_N/\pi_S$ | | | | | | | | 0.09 | 0.59 | -0.74** |
| Gen. time | | | | | | | | | 0.22 | -0.13 |
| $u$ | | | | | | | | | | -0.92** |

Asterisks indicate strength of support (**: $pp > 0.975$, *: $pp > 0.95$)

Table S2. Correlation coefficients between $dS$, $dN/dS$, $\pi_S$, $\pi_N/\pi_S$, life-history traits and $N_e$ using the estimates of $\pi_S$ and $\pi_N/\pi_S$ without hyper-geometric sampling

| | $dS$ | $dN/dS$ | Mat. | Mass | Long. | $\pi_S$ | $\pi_N/\pi_S$ | Gen. | $u$ | $N_e$ |
|---|---|---|---|---|---|---|---|---|---|---|
| $dS$ | | 0.26 | -0.38 | -0.65** | -0.28 | -0.61* | 0.47 | -0.43 | 0.71** | -0.72** |
| $dN/dS$ | | | 0.15 | 0.06 | 0.49 | -0.53 | 0.47 | 0.42 | 0.60* | -0.61* |
| Maturity | | | | 0.61** | 0.52** | 0.06 | 0.14 | 0.64** | 0.09 | -0.01 |
| Mass | | | | | 0.53** | 0.38 | -0.19 | 0.64** | -0.19 | 0.33 |
| Longevity | | | | | | -0.23 | 0.26 | 0.88** | 0.36 | -0.31 |
| $\pi_S$ | | | | | | | -0.79** | -0.03 | -0.67* | 0.92** |
| $\pi_N/\pi_S$ | | | | | | | | 0.15 | 0.62 | -0.77** |
| Gen. time | | | | | | | | | 0.30 | -0.16 |
| $u$ | | | | | | | | | | -0.89** |

Asterisks indicate strength of support (**: $pp > 0.975$, *: $pp > 0.95$)

Table S3. Estimates of mutation rate per year $r$ and per generation $u$ (posterior median and 96% credible interval), for extant and ancestral species, under a uniform prior over divergence times.

| species | $r$ (per $10^9$ years) | | $u$ (per $10^8$ generation) | | |
|---|---|---|---|---|---|
| | without anc. pol.[a] | with anc. pol.[b] | without anc. pol.[a] | with anc. pol.[b] | pedigrees[c] |
| *Homo* | 0.79 ( 0.59, 1.11) | 0.65 ( 0.48, 0.90) | 2.29 ( 1.70, 3.21) | 1.90 ( 1.38, 2.61) | 1.23 - 1.29 |
| *Pan* | 0.75 ( 0.61, 0.90) | 0.63 ( 0.50, 0.76) | 1.79 ( 1.48, 2.15) | 1.51 ( 1.21, 1.83) | 1.26 - 1.48 |
| *Gorilla* | 0.76 ( 0.43, 1.19) | 0.54 ( 0.29, 0.90) | 1.44 ( 0.82, 2.27) | 1.03 ( 0.56, 1.70) | 1.13 |
| *Pongo* | 0.86 ( 0.54, 1.23) | 0.73 ( 0.45, 1.06) | 2.14 ( 1.36, 3.08) | 1.83 ( 1.12, 2.65) | 1.66 |
| *Macaca* | 0.62 ( 0.48, 0.77) | 0.53 ( 0.40, 0.68) | 0.87 ( 0.68, 1.08) | 0.74 ( 0.56, 0.95) | 0.37 |
| *Papio* | 0.91 ( 0.50, 1.69) | 0.70 ( 0.37, 1.30) | 1.24 ( 0.73, 1.98) | 0.97 ( 0.54, 1.58) | 0.55 |
| *Aotus* | 0.96 ( 0.52, 1.67) | 0.91 ( 0.46, 1.61) | 1.03 ( 0.58, 1.65) | 0.96 ( 0.50, 1.56) | 0.81 |
| Catarrhini | 0.87 ( 0.52, 1.42) | 0.91 ( 0.54, 1.60) | 1.18 ( 0.78, 1.73) | 1.20 ( 0.76, 1.86) | |
| Platyrrhini | 1.49 ( 1.01, 2.21) | 1.43 ( 0.93, 2.18) | 1.56 ( 1.14, 2.15) | 1.50 ( 1.06, 2.12) | |
| Haplorrhini | 2.77 ( 1.33, 5.76) | 2.97 ( 1.44, 6.46) | 1.78 ( 1.03, 3.09) | 2.05 ( 1.16, 3.68) | |
| Strepsirrhini | 3.42 ( 1.64, 7.04) | 3.80 ( 1.81, 8.45) | 1.98 ( 1.16, 3.48) | 2.33 ( 1.32, 4.34) | |
| Primates | 3.04 ( 1.47, 6.39) | 3.22 ( 1.54, 7.08) | 1.86 ( 1.09, 3.29) | 2.10 ( 1.18, 3.86) | |

[a] naive-phylogenetic method (not accounting for ancestral polymorphism); [b] mean-coalescent method (accounting for ancestral polymorphism); [c] from Table 1 of Wu et al (2019)

Table S4. Estimates of effective population size ($\times 10^{-3}$, posterior median and 95% credible interval) for extant taxa and ancestors, under a uniform prior over divergence times.

| species | mechanistic | mech. w/o anc. pol. | phenomenological | uncoupled | coal.[a] | hmmcoal[b] |
|---|---|---|---|---|---|---|
| *Homo* | 23 ( 17, 32) | 19 ( 14, 26) | 19 ( 12, 35) | 21 ( 13, 35) | (13, 16) | 8 |
| *Pan* | 69 ( 57, 86) | 58 ( 48, 70) | 72 ( 45, 114) | 61 ( 39, 103) | (31, 62) | 30 |
| *Gorilla* | 69 ( 24, 184) | 67 ( 26, 177) | 102 ( 34, 334) | 48 ( 20, 117) | (28, 57) | 21 |
| *Pongo* | 42 ( 14, 113) | 37 ( 13, 93) | 50 ( 19, 125) | 32 ( 10, 106) | (42, 85) | 19 |
| *Homo-Pan* | 46 ( 28, 74) | 45 ( 28, 72) | 49 ( 29, 88) | 40 ( 24, 65) | (10, 47) | 50 |
| *Homo-Gorilla* | 47 ( 26, 78) | 47 ( 28, 80) | 53 ( 29, 98) | 41 ( 24, 71) | (27, 61) | 47 |
| Hominidae | 53 ( 26, 106) | 45 ( 23, 86) | 53 ( 27, 111) | 48 ( 22, 105) | | |
| Hominoidea | 62 ( 30, 131) | 57 ( 29, 112) | 70 ( 34, 159) | 58 ( 25, 137) | | |
| Cercopithecidae | 67 ( 35, 125) | 74 ( 42, 128) | 79 ( 41, 156) | 73 ( 37, 152) | | |
| Catarrhini | 74 ( 35, 156) | 73 ( 38, 145) | 69 ( 33, 155) | 58 ( 26, 131) | | |
| Platyrrhini | 42 ( 21, 83) | 37 ( 20, 71) | 32 ( 16, 61) | 31 ( 13, 75) | | |
| Simiiformes | 60 ( 26, 138) | 56 ( 27, 124) | 37 ( 15, 87) | 36 ( 14, 90) | | |
| *Tarsius* | 536 ( 99, 4855) | 407 ( 88, 2558) | 107 ( 22, 561) | 56 ( 5, 599) | | |
| Haplorrhini | 92 ( 37, 239) | 83 ( 36, 207) | 32 ( 9, 97) | 34 ( 11, 103) | | |
| Lorisiformes | 125 ( 58, 279) | 124 ( 61, 286) | 46 ( 14, 120) | 49 ( 20, 121) | | |
| Lemuriformes | 94 ( 41, 202) | 90 ( 41, 187) | 89 ( 34, 218) | 123 ( 51, 310) | | |
| *Daubentonia* | 1251 ( 208, 11831) | 1296 ( 237, 8859) | 1030 ( 181, 8983) | 443 ( 56, 3733) | | |
| Strepsirrhini | 91 ( 35, 246) | 82 ( 34, 214) | 27 ( 7, 91) | 33 ( 12, 92) | | |
| Primates | 93 ( 35, 240) | 80 ( 34, 201) | 29 ( 7, 94) | 32 ( 11, 97) | | |

[a] from Prado-Martinez et al, 2013, table 1, for extant hominids, and from Rannala and Yang, 2003 for ancestral species; [b] from Prado-Martinez et al, 2013, figure 2

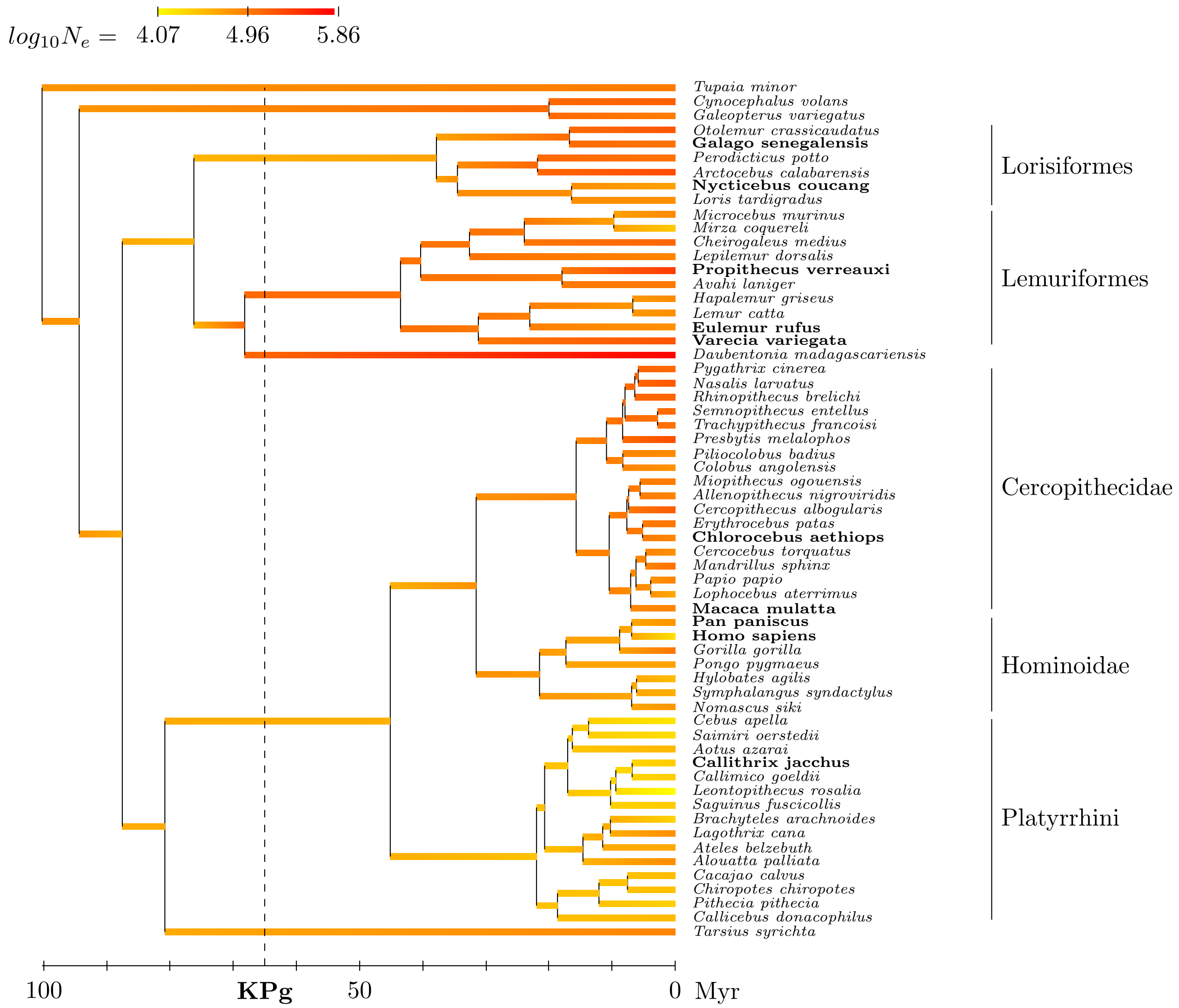


Figure S1. Reconstructed phylogenetic history of $N_e$ (posterior median estimate) under the phenomenological model.

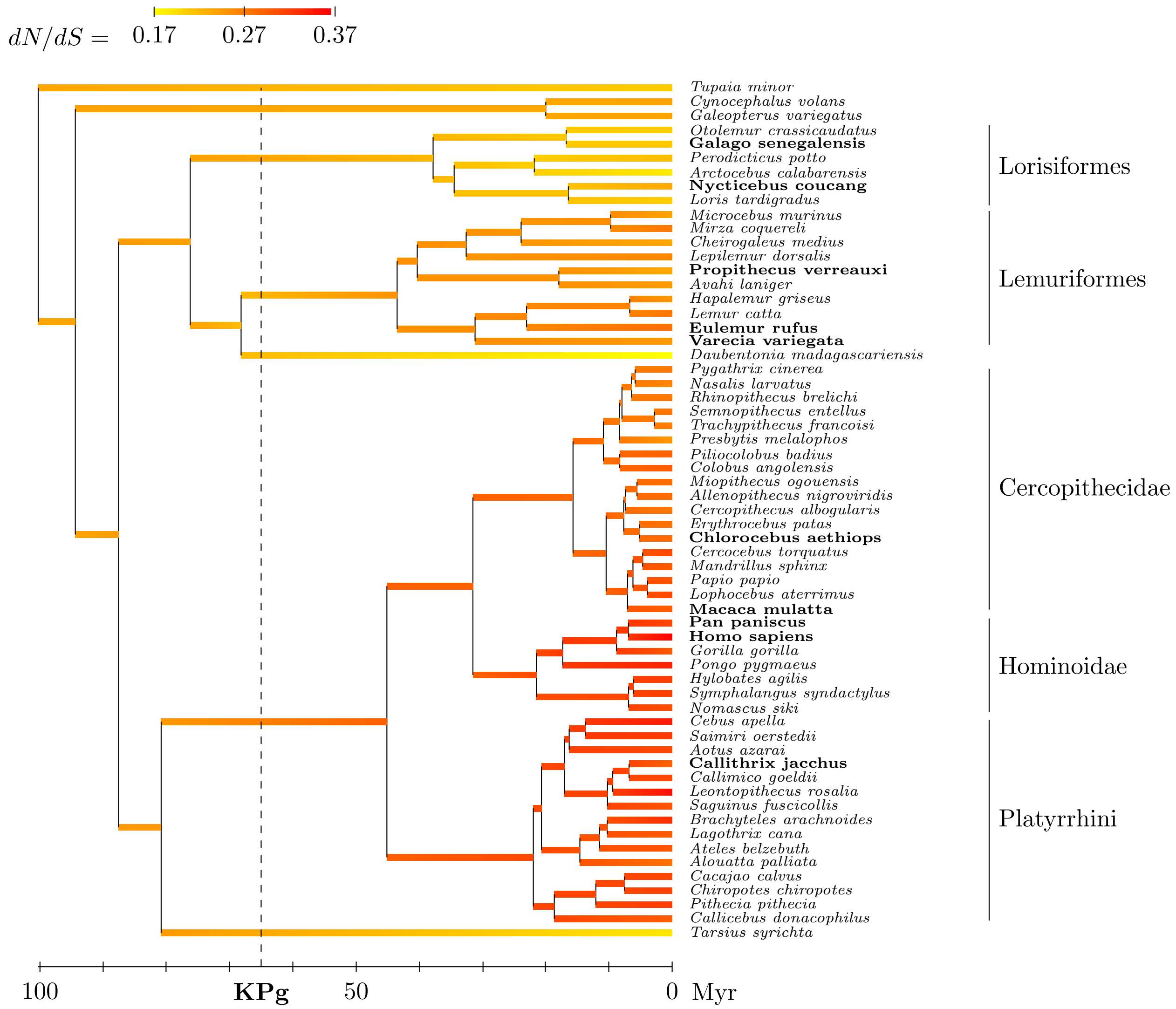


Figure S2. Reconstructed phylogenetic history of $dN/dS$ (posterior median estimate) under the phenomenological model.

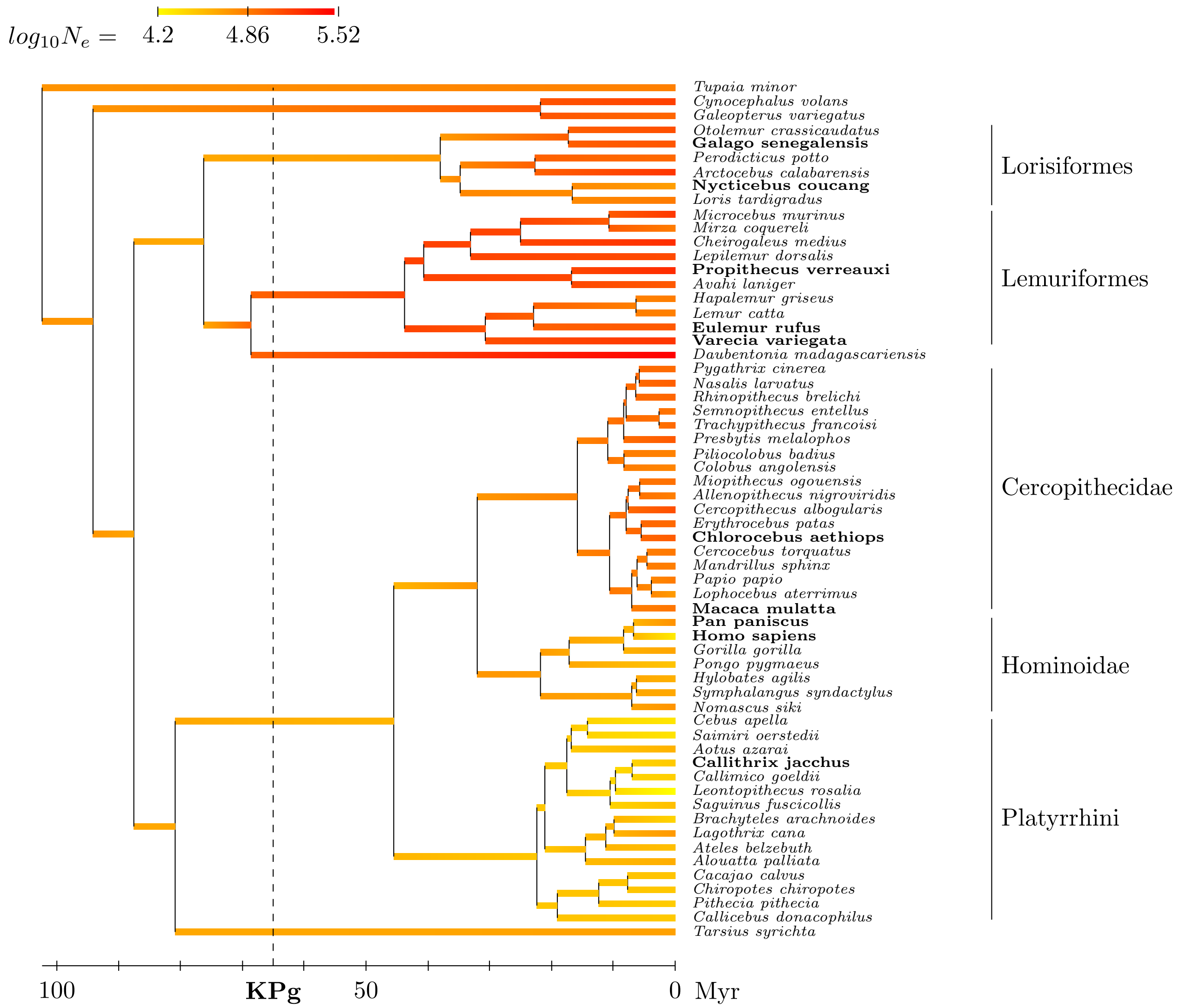


Figure S3. Reconstructed phylogenetic history of $N_e$ (posterior median estimate) under the uncoupled model.

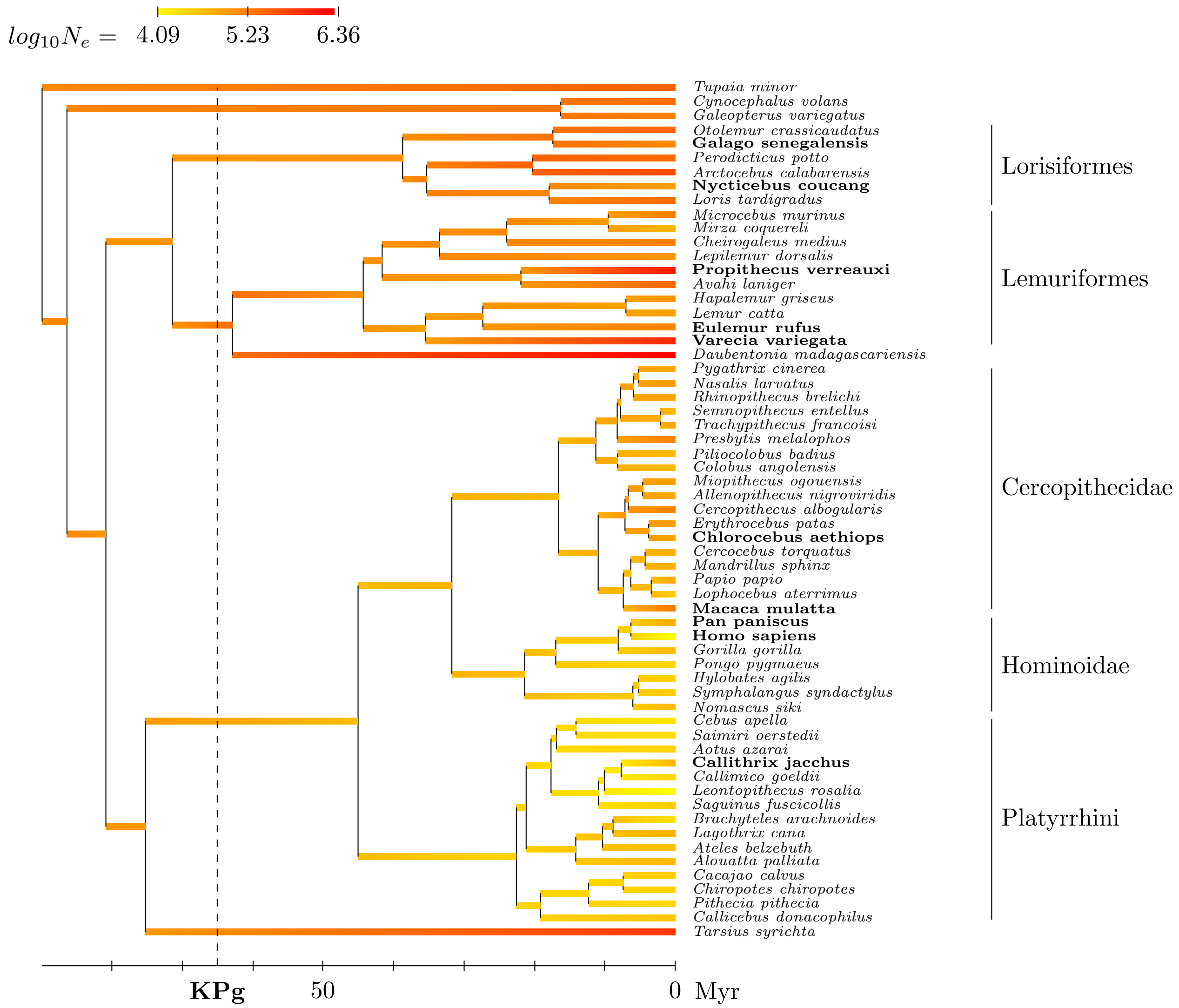


Figure S4. Reconstructed phylogenetic history of $N_e$ under the mechanistic model, with $\beta$ fixed at 0.16.

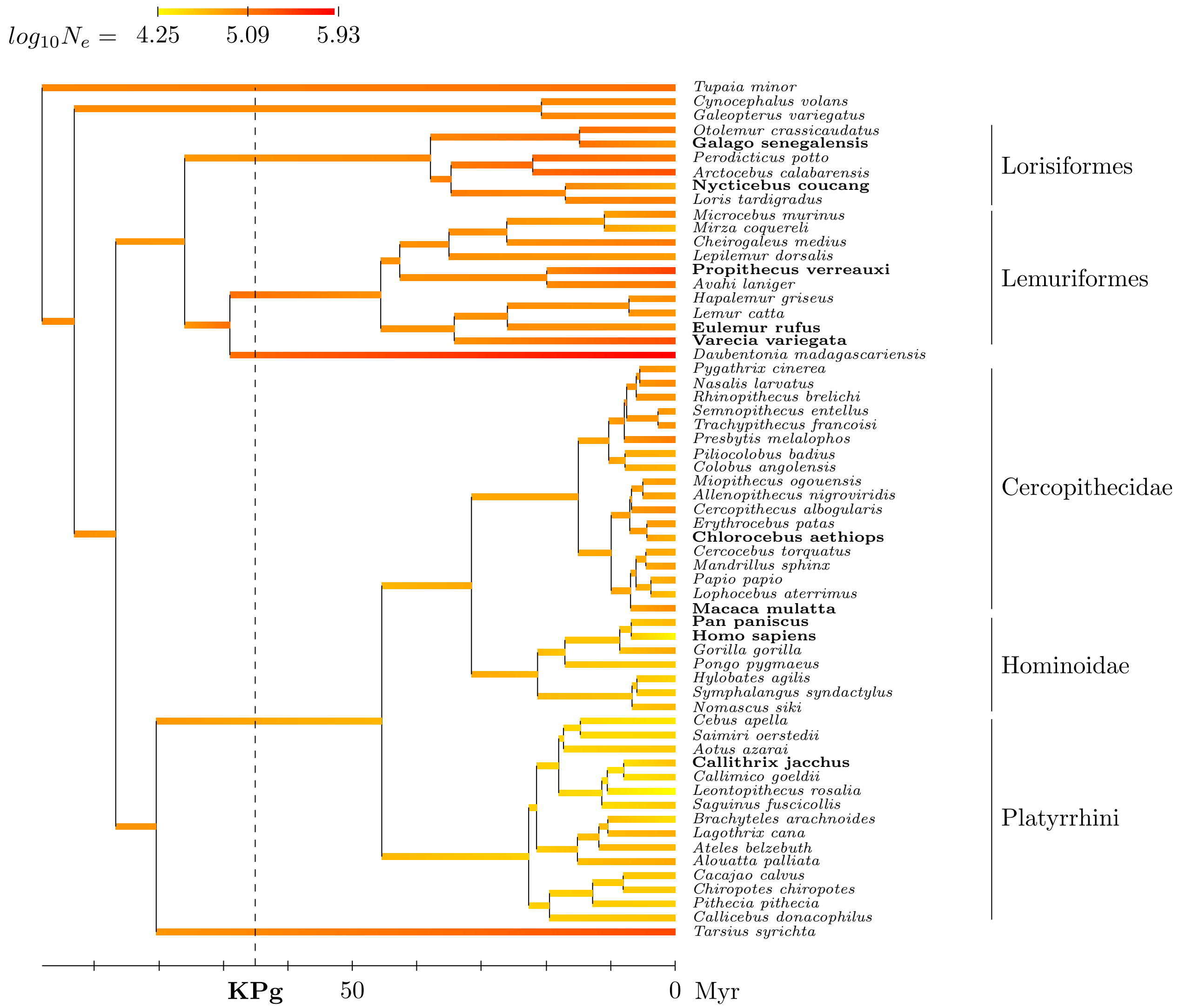


Figure S5. Reconstructed phylogenetic history of $N_e$ (posterior median estimate) under the mechanistic model, without ancestral polymorphism.

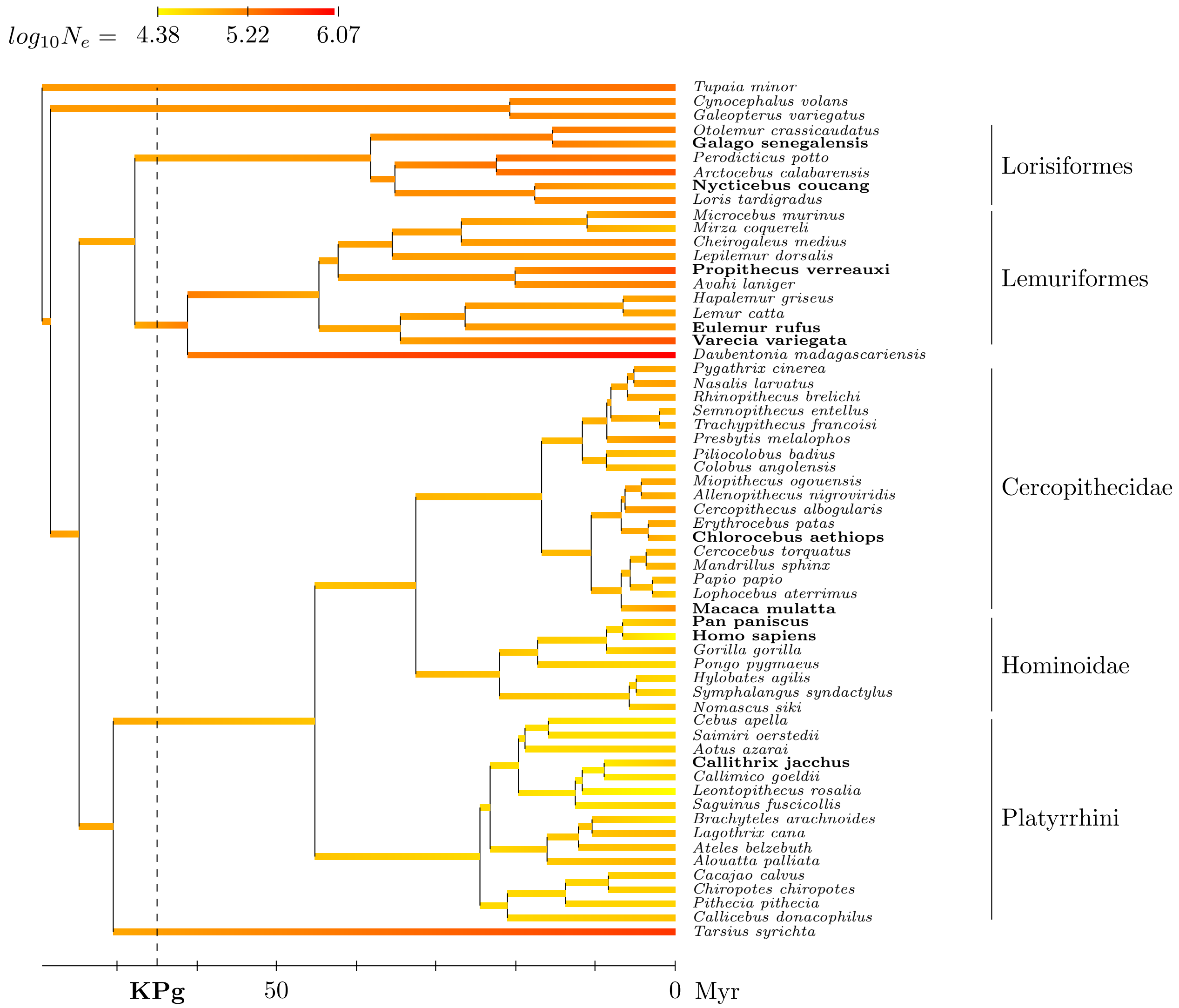


Figure S6. Reconstructed phylogenetic history of $N_e$ (posterior median estimate) under the mechanistic model and using a uniform prior over divergence times.

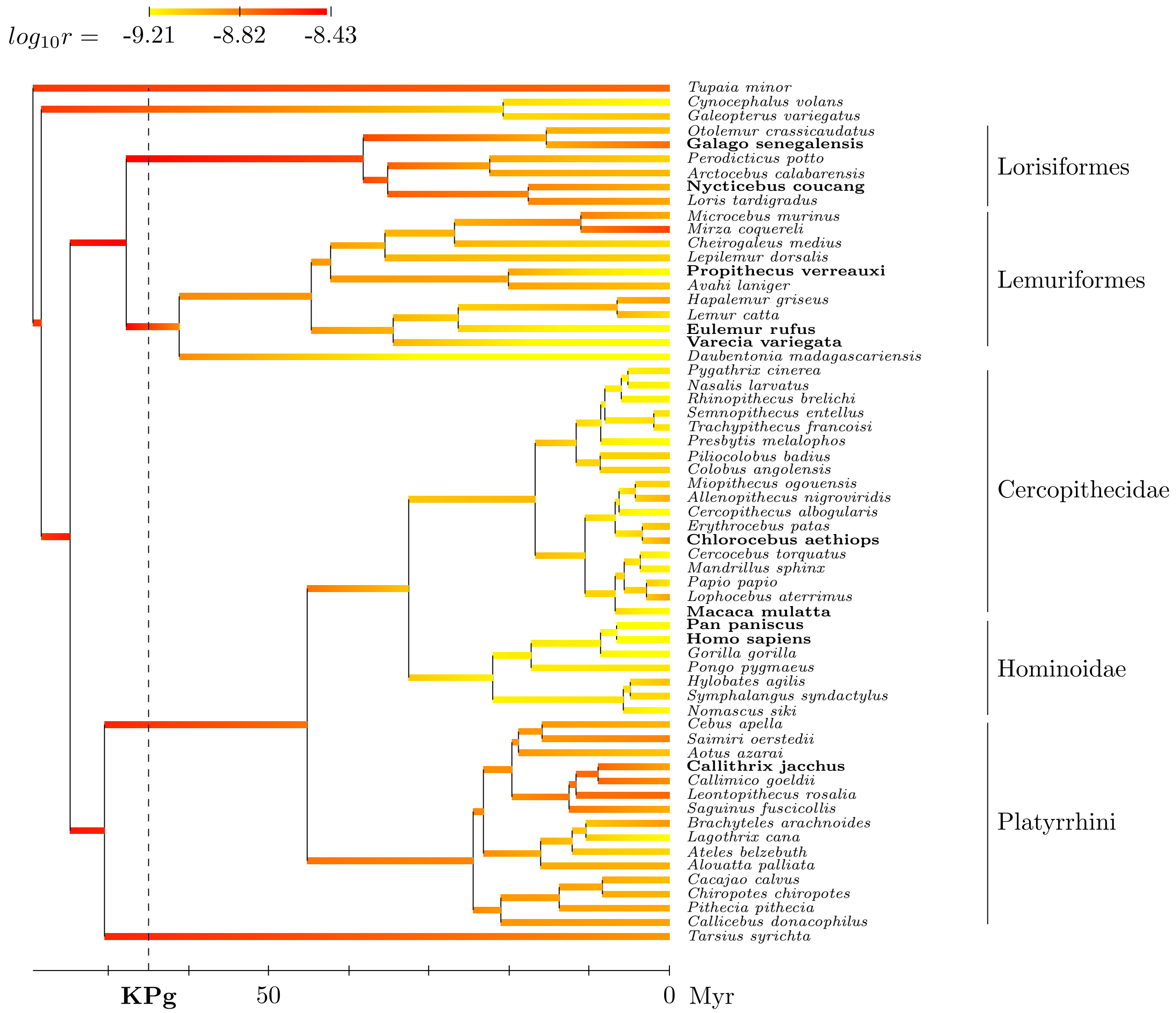


Figure S7. Reconstructed phylogenetic history of $r$ (posterior median estimate) under the mechanistic model and using a uniform prior over divergence times.

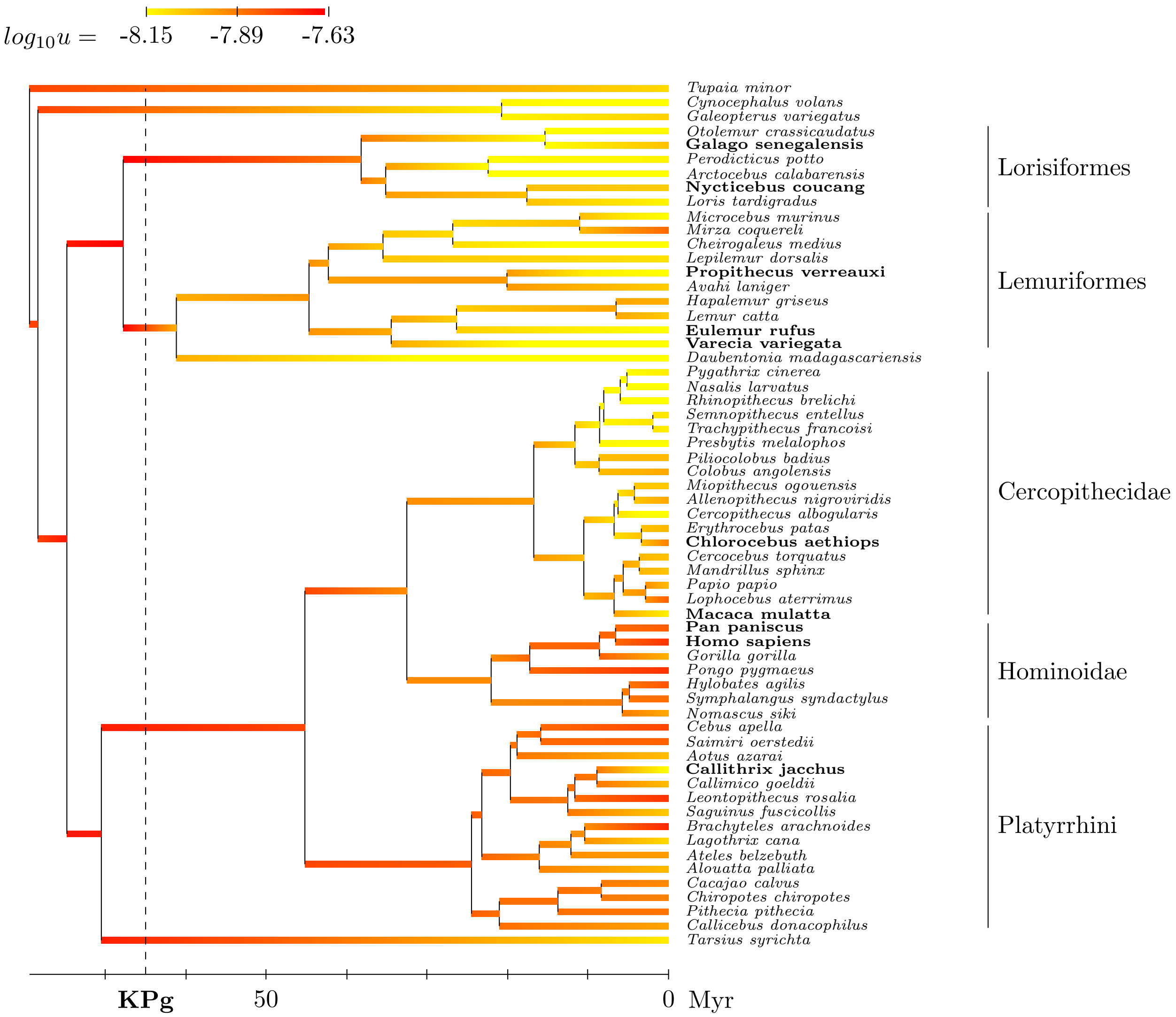


Figure S8. Reconstructed phylogenetic history of $u$ (posterior median estimate) under the mechanistic model and using a uniform prior over divergence times.
